# Amoeba predation of *Cryptococcus*: A quantitative and population genomic evaluation of the Accidental Pathogen hypothesis

**DOI:** 10.1101/2022.12.08.519367

**Authors:** Thomas J. C. Sauters, Cullen Roth, Debra Murray, Sheng Sun, Anna Floyd-Averette, Chinaemerem U. Onyishi, Robin C. May, Joseph Heitman, Paul M. Magwene

## Abstract

The “Amoeboid Predator-Fungal Animal Virulence Hypothesis” posits that interactions with environmental phagocytes shape the evolution of virulence traits in fungal pathogens. In this hypothesis, selection to avoid predation by amoeba inadvertently selects for traits that contribute to fungal escape from phagocytic immune cells. Here, we investigate this hypothesis in the human fungal pathogens *Cryptococcus neoformans* and *Cryptococcus deneoformans*. Applying quantitative trait locus (QTL) mapping and comparative genomics, we discovered a cross-species QTL region that is responsible for variation in resistance to amoeba predation. In *C. neoformans*, this same QTL was found to have pleiotropic effects on melanization, an established virulence factor. Through fine mapping and population genomic comparisons, we identified the gene encoding the transcription factor Bzp4 that underlies this pleiotropic QTL and we show that decreased expression of this gene reduces melanization and increases susceptibility to amoeba predation. Despite the joint effects of *BZP4* on amoeba resistance and melanin production, we find no relationship between *BZP4* genotype and escape from macrophages or virulence in murine models of disease. Our findings provide new perspectives on how microbial ecology shapes the genetic architecture of fungal virulence, and suggests the need for more nuanced models for the evolution of pathogenesis that account for the complexities of both microbe-microbe and microbe-host interactions.

**Author summary:** A prominent hypothesis for the evolution of many environmental pathogens proposes that opportunistic pathogenesis is an “accidental” by-product of selection to survive encounters with microbial predators. Chief among the predators that have been suggested as relevant to the evolution of virulence are phagocytic amoebae. Amoebae share many characteristics with macrophages and other primary immune cells that microbial pathogens encounter during infection of animal hosts. This has led to the suggestion that amoebae may act as “training grounds” for both bacterial and fungal pathogens. In this study we test key tenets of the accidental pathogen hypothesis by examining two related questions: “Do alleles important for survival in the face of amoeba predation correspond to known virulence genes? And does genetic variation that increases resistance to amoeba predation increase virulence potential?” We carried out quantitative trait locus (QTL) mapping in two species of the human fungal pathogen *Cryptococcus* and identified an orthologous QTL, shared by the two species, where allelic variation is a key predictor of resistance to amoeba predation. In *C. neoformans* we show that this QTL corresponds to a deletion upstream of a transcription factor gene, *BZP4*. Variation at *BZP4* also predicts melanin synthesis, another trait implicated in *Cryptococcus* virulence. Although *BZP4* genotype is a strong predictor of resistance to amoeba predation, we find no correlation between genetic variation at this locus and the ability to proliferate in macrophages or to kill animal hosts. Our findings suggest that the evolutionary landscape of fungal virulence is complex, and highlights the importance of accounting for natural genetic variation when evaluating evolutionary hypotheses.

## Introduction

For many free-living pathogens, there is no host-to-host transmission and infection of a host is not an obligatory stage of their life cycle. Pathogenesis in these cases is considered opportunistic, and key traits that facilitate virulence are not likely to have evolved due to adaptation to the host directly [1, 2]. Rather, the ability to cause disease is hypothesized to be an unintentional byproduct of evolving in a varied, stressful environment (“accidental virulence”; [3]). This raises the question: “What environmental interactions contribute to the evolution of virulence?”

A prominent hypothesis proposed for many environmental pathogens suggests that predator-prey interactions between microbes drive the evolution of traits advantageous to pathogenesis [4]. Chief among the predators that have been suggested as relevant to the evolution of virulence traits are amoebae. For example, bacterial pathogens such as *Bordetella*, *Legionella*, and *Pseudomonas*, as well as the fungal pathogens *Paracoccidioides*, *Cryptococcus*, and *Aspergillus*, are all preyed upon by phagocytic amoebae [5–11]. Amoebae share many similarities with macrophages and other primary immune cells that microbial pathogens encounter during infection of mammalian hosts. These similarities include immune receptors that detect microbial PAMPs, actin mediated phagocytosis, acidification, nitrosative stress, and metalo-ion toxicity in the phagosome. [12–15]. In light of this, it has been proposed that amoebae may serve as “training grounds” for intracellular pathogens [16]. For fungi in particular, the idea that interactions with amoebae in the environment drives the selection of fungal traits necessary for survival during mammalian infection has been termed the “Amoeboid Predator-Fungal Animal Virulence Hypothesis” [17].

For the fungal pathogen *Cryptococcus neoformans*, interactions with free-living amoebae have been documented for nearly 100 years [18]. Amoebae are found in many of the same niches that *C. neoformans* inhabits, and *C. neoformans* is actively consumed by amoebae isolated from pigeon guano [19]. *C. neoformans*, and the amoebae that consume it, are globally distributed [20, 21]. *C. neoformans* is a saprophytic fungus; however, it has the ability to cause disease in vulnerable human populations, primarily infecting individuals with reduced immunity due to factors such as HIV/AIDS or immunosuppressive drug treatments [22–25]. *C. neoformans* infections in mammals are facilitated by a variety of traits including: a polysaccharide capsule [26–28], the ability to grow at high temperatures [29, 30], the production of melanin [31, 32], and a battery of secreted phospholipases [33, 34] and ureases [35, 36]. These same virulence factors act as defense mechanisms against amoebae [28, 37, 38]. Passaging *C. neoformans* strains with amoebae increases virulence factor presentation and results in enhanced pathogenicity in mammalian tissue culture, insects, and mouse models of infection [39, 40]. These findings support the hypothesis that amoebae may play a key role in the evolution of *C. neoformans* virulence factors. However, most studies that characterize the similarities between *Cryptococcus*’s interactions with amoebae and with animal immune systems have targeted known virulence genes, primarily through gene deletion studies. Furthermore, these studies analyzed a small number of *Cryptococcus* strain backgrounds. Focusing on previously identified genes, in a limited number of strains, may bias or obscure other genes and pathways important for amoeba resistance.

In this study we ask, “Do alleles important for fungal survival with amoeba correspond to known virulence genes?” To answer this question, we employed quantitative trait locus (QTL) mapping to identify genomic regions and allelic variants that contribute to resistance against amoeba predation in two pathogenic species of *Cryptococcus*, *C. neoformans* and *C. deneoformans*. For both species we identified major effect QTL. Surprisingly, these QTL regions were found to be homologous between the species. For *C. neoformans*, the amoeba resistance QTL identified is also a melanization QTL. By combining comparative genomics and genetic engineering we identified a likely causal variant for this QTL region, a 1.8 kb deletion upstream of the transcription factor encoding gene *BZP4*. Disruption of this region leads to altered transcription of *BZP4* and other genes, and these transcriptional differences are in turn associated with reduced amoeba resistance and melanization capacity. Despite alterations in amoeba resistance and melanization associated with mutation of *BZP4*, comparative analysis suggests that *BZP4* is not required for virulence in mice or macrophages. In addition, no relationship is found between genetic variation in the ability to resist amoeba predation and virulence in mouse models of infection. Our findings not only advance the understanding of the genetic architecture of virulence traits, but also suggest the need for a more nuanced perspective on the evolutionary and ecological interactions that have shaped microbial pathogenesis.

## Materials and methods

### Strains, Laboratory Crosses and Isolation

All strains were maintained on yeast peptone dextrose (YPD) plates grown at 30*^◦^*C for 48 hours from −80*^◦^*C stocks. Overnight cultures for amoeba assays, melanin assays, and RNA isolation were made in liquid YPD at 30*^◦^*C on a rotor drum.

### Amoeba Resistance Assay

*C. neoformans* and *C. deneoformans* strains were grown overnight in 3 mL of liquid YPD on a roller drum before being diluted down to OD_600_ 0.6. 100 µL of diluted culture was spread on solid V8 media petri dishes using glass beads. Plates were grown at 30*^◦^*C for 60 hours before being removed from the incubator. *Acanthamoeba castellanii* (ATCC 30234) were grown in ATCC 712 in a 75 mL tissue culture flask. Amoeba were harvested, between passage 5 and 15, from flasks and suspended at a concentration of 10^6^ cells/mL before 50 µL of amoeba culture were pipetted onto the center of the *Cryptococcus* lawn. Plates were allowed to dry at room temperature on the bench top for 10 minutes and then placed in a 25*^◦^*C incubator for 12 - 18 days. Measurements were taken at days 1, 12, and 18. Area of clearance was calculated by subtracting the day 1 measurement from the final day measurement as the day 1 measurement represented the initial spread of the amoeba culture on the plates. Two final time points were used (12 and 18 days) based on amoeba replication rate and activity. Rank order of amoeba affect was conserved between days 12 and 18.

### Melanization assay

Melanization was assayed by growing strains on minimal media plates with L-DOPA (7.6 mM L-asparagine monohydrate, 5.6 mM glucose, 10 mM MgSO4, 0.5 mM 3,4-dihydroxy-L-phenylalanine, 0.3 mM thiamine-HCl, and 20 nM biotin) for 72 hours. Plates were then scanned on an Epson Expression XL Flatbed Scanner in reflective mode at 300 dpi. ImageJ was used to calculate greyscale intensity of the colonies. Each sample was measured in triplicate.

### Spore Dissection

Meiotic progeny were recovered by microdissection of random basidiospores as previously described [81]. Briefly, cells from the two parental strains were each resuspended in sterile water to a density of OD_600_=1.0. Equal volumes of cell suspensions were mixed, and 5 *µ*L of the mixture, as well as the two parental strains (serving as negative controls of mating), were spotted onto MS solid medium. The MS plates were incubated in the dark at room temperature (23*^◦^*C for two weeks, at which time robust hyphae, basidia, and basidiospore chains were produced by the spots from the mixture of the two parental strains. Basidiospores from a large number of basidia in one location along the edge of the mating spot were picked directly from the MS plates using the needle of a dissection microscope each time, then transferred and separated onto YPD solid medium. To reduce the chances of sampling clones from the same basidia, we only separated limited numbers of basidiospores from one location (<5%), and sampled multiple locations, as well as from multiple mating spots.

### DNA extraction, Library Preparation, and Sequencing

DNA was extracted with MasterPure Yeast DNA Purification kit and cleaned up the Zymo Research Genomic Clean and Concentrator kit (following manufacturer’s instructions) followed by quantification with PicoGreen. After quantifying the DNA with PicoGreen, samples were prepped for genomic sequencing using seqWell’s plexWell 96 kit to prepare the libraries. Briefly, samples were individually bar-coded in sets of 96 using randomly inserted transposons, pooled, and then purified. Next, each pooled sample was bar-coded, enriched, and finally size-selected purified. Libraries were sequenced at Duke University’s Sequencing and Genomic Technologies Facility on the NovaSeq 6000 S-Prime with 150 basepair paired end reads. Reads were aligned to the H99 *C. neoformans* reference genome using BWA. Variant calling was carried out using SAMtools and Freebayes.

### Segregants Filtering and SNP Filtering

Segregants were filtered to remove aneuploidy and clonality described in detail in [82]. Of the original 384 segregants we were left with 304 after filtering. Variant sites were filtered based on read depth, allelic read depth ratio, quality scores, and minor allele frequency as described in [47]. Total number of bi-allelic variant sites prior to filtering was 59,430 that were reduced down to 46,670 after filtering.

### QTL Mapping

The 46,670 genetic variants were combined into 4,943 haploblocks, defined by linkage, using methods described previously by Roth *et al.* [47]. For association testing of amoeba resistance and melanization, a Mann-Whitney U test was used across these 4,943 haploblocks to associate phenotype and genotype, coding the Bt22 and Ftc555-1 genotypes as zero and one respectively. The -log10 (p-value) from the Mann-Whitney U tests was monitored to identify QTL and 95% confidence intervals were calculated using permutation testing, a thousand times with replacement [83].

### Permutation Testing

Permutation testing was carried out as described in [84] and [47] for establishing significance thresholds for QTL mapping. A thousand permutations were used for the melanin and amoeba phenotypes. Random assignments of genotype and phenotpe were held constant for every condition tested to preserve autocorrelation between phenotypes. The 95*^th^* percentile of the permuted null distribution were used as the threshold for significance.

### Population Genomics

Raw Illumina sequencing reads generated by Desjardins et al. [48] were downloaded from the NIH Sequence Read Archive (BioProject ID PRJNA382844). For each of the 387 BioSamples (corresponding strains of interest) associated with the BioProject, we created a reference-based genome assembly based on aligning paired-end sequence data to the genome of the *C. neoformans* reference strain H99 (FungiDB R53). In cases where there were multiple sequencing runs for a given BioSample, we used the sequencing run containing the largest number of paired-end reads. To create reference-based genome assemblies we aligned reads to the H99 reference genome using BWA (v0.7.17-r1188; [85]), called variants using FreeBayes (v1.3.5; [86]), and generated strain-specific consensus assemblies by instantiating the called variants onto the reference genome. The read alignment, variant calling, and consensus assembly were carried out using the Snippy (https://github.com/tseemann/snippy) pipeline tool.

Following construction of consensus assemblies, genome feature annotation was “lifted over” from the H99 reference genome to each strain-specific genome using the software tool Liftoff (v1.6.3; [87]). The polish option of Liftoff was employed to re-align exons in cases where the lift-over procedure resulted in start/stop codon loss or introduced an in-frame stop codon. Based on the polished lift-over annotation, the AGAT GTF/GFF Toolkit software (https://github.com/NBISweden/AGAT) was used to predict protein sequences for all annotated genes in each strain-specific assembly using the agat_sp_extract_sequences.pl script. Where multiple protein isoforms are annotated in the reference genome, we generated predictions for each isoform.

Candidate *BZP4* loss-of-function alleles were identified as those cases where the predicted length of the amino acid sequence of Bzp4 is < 90% of the modal protein length estimated from the entire set of strains.

To identify candidate regulatory alleles upstream of *BZP4*, the SAMtools coverage program was employed to summarize the coverage and read depth in the 1 kb region upstream of *BZP4*. Strains where the proportion of covered bases was less than 80% and there was reduced read-depth relative to the surrounding genomic region were classified as candidate regulatory deletions. These candidate regions were subsequently confirmed by manual inspection of read alignments.

Population genetic sequence diversity statistics such as *π* and Tajima’s D were estimated for *BZP4* as well as all predicted protein coding genes on chromosome 8 and all 178 transcription factors identified by Jung et al. [58]. These estimates were calculated from multiple-sequence alignments generated from the reference based assemblies described above. Alignment were trimmed using ClipKIT [88], and the statistics were calculated using the population genetic statistical functions implemented in DendroPy [89].

### RNA Isolation and Sequencing

12 segregants and duplicates of the parental strains were used for the analyses comprising 16 individual samples. Samples were grown overnight in liquid YPD on a rollerdrum and were added to V8 petri dishes. V8 cultures were grown at 30*^◦^*C for 60 hours. Amoeba cultures were collected and suspended at a concentration of 1 *×* 10^6^ cells/mL. 450 µL of amoeba culture was added to the center of the *Cryptococcus* lawn with slight agitation to aid in the spread of the culture. Plates were allowed to dry on the benchtop for 30 minutes before being incubated at 25*^◦^*C for 48 hours. A consistent area of 30 cm^2^ was cut from the plates and then scraped to collect cells. Collected cells were resuspended in 1 mL of PBS and were placed in dry ice for 10 minutes. Samples were then lyophilized for 12 - 18 hours. Whole RNA was extracted using the RNAeasy Plant Mini Kit (Qiagen 74904).

Control samples did not have amoeba added to them, but were handled in a similar fashion in every other aspect of the protocol.

Libraries were prepared and sequenced by the Duke sequencing core using the Illumina NextSeq 500 High Output Kit producing 150-base pair paired end reads.

### RNAseq Analysis

Reads were aligned using the CNA3 of H99 *C. neoformans* var. grubii (accession GCA_000149245.3). from the Ensemble Fungi database. Reads were aligned using Kallisto.

Analysis of RNA sequences was performed using Deseq2 in R. Briefly, transcript abundance was normalized using a built in median of ratios method. Samples were normalized based on condition (amobea or control). GO term analysis was performed using the fungidb GeneByLocusTag tool. Correlation of gene expression with *BZP4* was performed using the normalized count output from Deseq2. The z-score for each gene was calculated before a Pearson correlation between *BZP4* and all other genes was established. Genes with a correlation of r > |0.5| were further subset by differential expression based on the *BZP4*4 allele.

### Sequence Motif Analysis

DNA sequence motif analysis of the upstream 1 kb regions of genes whose expression correlates with *BZP4* were carried out using the program XSTREME [54], part of the MEME Suite [55]. A control set of sequences was generated by randomly selecting the upstream regulatory sequences of 500 genes. An E-value of 10*^−^*8 was used as a cutoff to identify enriched sequence motifs. *BZP4*-correlated regulatory sequences were compared to three independent sets of control sequences, and we focused on motifs that were below the E-value cutoff in each of the three comparisons.

### Tissue Culture

The J774A.1 macrophage cell line was cultured in T-75 flasks [Fisher Scientific] in Dulbecco’s Modified Eagle medium, low glucose (DMEM) [Sigma-Aldrich], supplemented with 10% live fetal bovine serum (FBS) [Sigma-Aldrich], 2mM L-glutamine [Sigma-Aldrich], and 1% Penicillin and Streptomycin solution [Sigma-Aldrich] at 37*^◦^*C and 5% CO^2^.

### Phagocytosis Assay

To measure the phagocytosis of various *Cryptococcus* segregants by macrophages, J774A.1 cells were seeded at a density of 1×10^5^ cells per well of a 24-well plate [Greiner Bio-One], then incubated overnight at 37*^◦^*C and 5% CO^2^. At the same time, an overnight culture of *C. neoformans* parental strains or segregants was set up by picking a fungal colony from YPD agar plates (50g/L YPD broth powder [Sigma-Aldrich], 2% Agar [MP Biomedical]) and resuspending it in 3mL liquid YPD broth (50 g/L YPD broth powder [Sigma-Aldrich]). The culture was then incubated at 25*^◦^*C overnight under constant rotation (200 rpm).

On the day of the assay, macrophages were activated using 150ng/mL phorbol 12-myristate 13-acetate (PMA) [Sigma-Aldrich] for 1 hour at 37*^◦^*C. PMA stimulation was performed in serum-free media to eliminate the contribution of complement proteins during phagocytosis. To prepare *C. neoformans* for infection, overnight *C. neoformans* cultures were washed two times in 1X PBS, counted using a hemacytometer, and fungi was incubated with macrophages at a multiplicity of infection (MOI) of 10:1. The infection was allowed to take place for 2h at 37 *^◦^*C and 5% CO^2^. After 2 h infection, as much extracellular *Cryptococcus* as possible was washed off using 1X PBS.

### Fluorescent Microscopy Imaging

The number of phagocytosed fungi was quantified from images from a fluorescent microscope. To distinguish between phagocytosed and extracellular *C. neoformans*, wells were treated with 10 μg/mL calcofluor white (CFW) [Sigma-Aldrich] for 10 mins at 37*^◦^*C. The wells were washed again with PBS to remove residual CFW. Fluorescent microscopy images were acquired at 20X magnification using the Nikon Eclipse Ti inverted microscope [Nikon] fitted with the QICAM Fast 1394 camera [Hamamatsu]. Images were analysed using the Fiji image processing software [ImageJ]. To quantify the number of phagocytosed *Cryptococcus* from the resulting images, the total number of ingested *C. neoformans* was counted in 200 macrophages, then the values were applied to the following equation: ((number of phagocytosed *C. neoformans*/number of macrophages) *×* 100).

### Intracellular Proliferation Rate Assay and Time-lapse Imaging

To investigate the intracellular proliferation rate (IPR) of *Cryptococcus* strains within macrophages, infected macrophages were captured at a regular interval over an extended period. Time-lapse imaging was performed by running the phagocytosis assay as usual, then after washing off extracellular *Cryptococcus* with 1X PBS, serum-free culture media was added back into the wells before imaging. Images were captured using the Nikon Eclipse Ti microscope at 20X magnification. Images were acquired every 5 minutes for 18 hours at 37 *^◦^*C and 5% CO2.

The resulting video was analysed using Fiji [ImageJ] and IPR was determined by quantifying the total number of internalised fungi in 200 macrophages at the ‘first frame’ (time point 0 (T0)) and ‘last frame’ (T10). The resulting values were used in the following equation: ((number of phagocytosed *C. neoformans*/number of macrophages) *×* 100). Next, the number of phagocytosed fungi at T10 was divided by the number of phagocytosed fungi at T0 to give the IPR (IPR = T10/T0).

### Mouse Infections

*Cryptococcus* strains for inoculation were grown overnight in 5 ml of YPD broth at 30*^◦^*C in a roller drum. Cells were pelleted by centrifugation and washed twice with sterile PBS. The cell pellet was resuspended in PBS, diluted, and counted by hemocytometer. The final inoculum was adjusted to a cell density of 4 x 10^6^ CFU/ml. Test groups consisting of five male and five female A/J mice aged 4-5 weeks were purchased from Jackson Labs (stock #000646) and infected via intranasal instillation. Mice were anesthetized using isoflurane administered with a calibrated vaporizer. 25 µl of the prepared inoculum was pipetted into the nares one drop at a time until the full volume containing 10^5^ CFU was inhaled. Mice were observed until fully recovered from anesthesia. Following infection, mice were monitored daily for symptoms of disease progression including weight loss, labored breathing, lack of grooming, social isolation, and any signs of pain or distress. Mice were euthanized upon reaching humane endpoints according to guidelines set forth by Duke University’s Animal Care and Use Program. Survival curves were plotted using GraphPad Prism version 8 and analyzed using log-rank (Mantel-Cox) statistical test.

### Ethics Statement

Animal experiments were performed under Duke protocol number A148-19-07, in accordance with guidance issued by Duke’s Institutional Animal Care and Use Committee and the U.S. Animal Welfare Act. Animals were housed in facilities managed by veterinary staff with Duke Lab Animal Research (DLAR) and accredited by the Association for Assessment and Accreditation of Laboratory Animal Care (AAALAC).

### Data Availability

Genome sequence rdata for the Bt22 *×* Ftc555-1 mapping population are available from the NIH Sequence Read Archive (BioProject ID PRJNA932005). RNA-seq data are available from the NIH Gene Expression Omnibus (Series GSE238170).

All code and data used to generate the figures in this paper are available at https://github.com/magwenelab/amoeba-qtl-code. RNAseq, virulence, and basic amoeba survival curve figures were all generated using R. QTL analysis and related figures were generated in Python.

An overview of the strains used in each experiment is included in Table S5.

## Results

### Comparison of Amoeba Resistance in Diverse *C. neoformans* Strains

We developed a plate-based assay to quantify *Cryptococcus* resistance to predation by the amoeba, *Acanthamoeba castellanii* (Fig 1A; [41]). Briefly, an established lawn of *Cryptococcus* cells was inoculated with a drop of amoeba. After a defined period of time, the cleared (consumed) portion of the lawn is quantified and used as a measure of resistance. The larger the clearance area, the less resistant the cells are to amoeba consumption.

**Fig 1.**
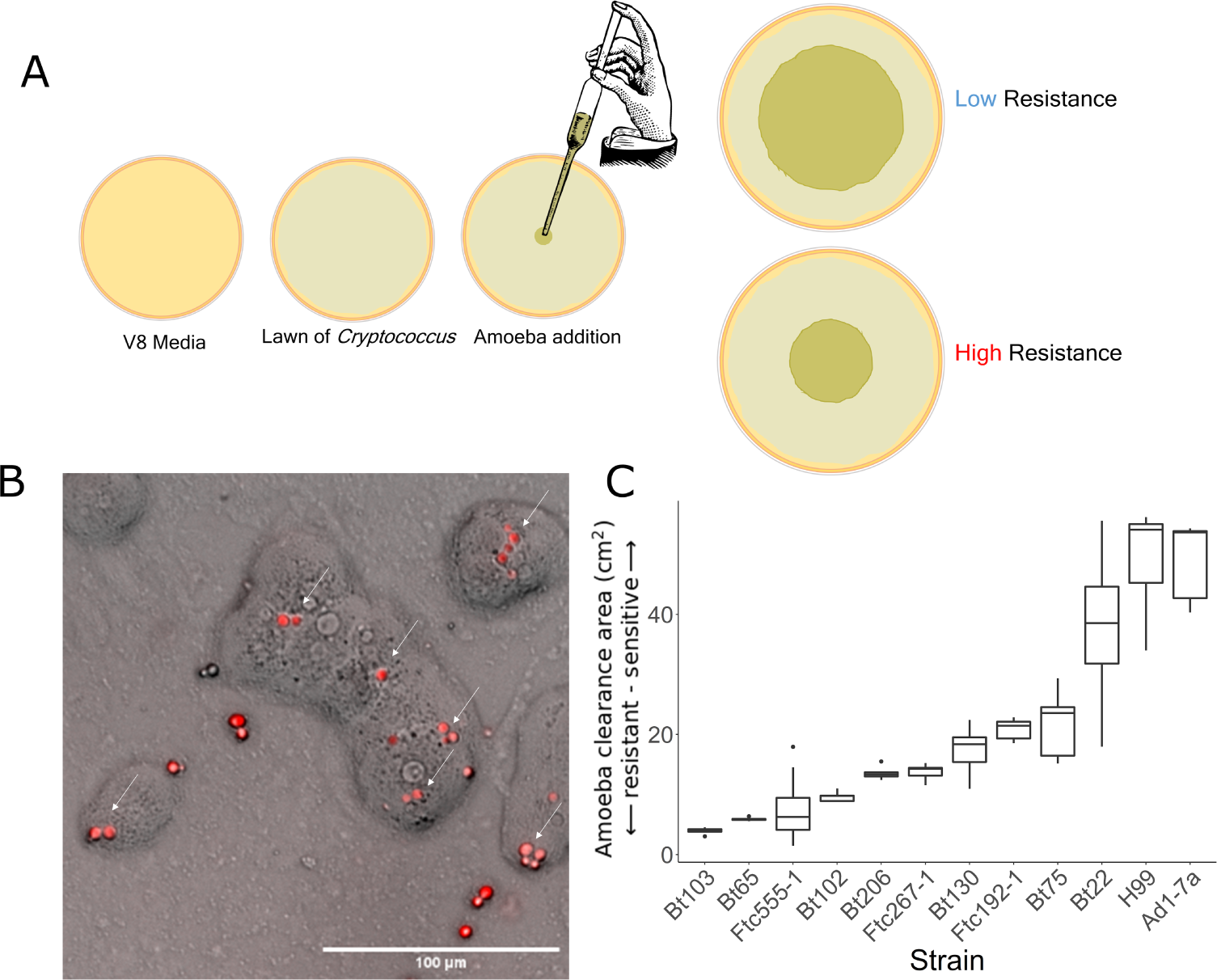
Amoeba resistance varies between strains of *C. neoformans*. **A.** A schematic overview of the amoeba resistance assay. A lawn of *Cryptococcus* is grown on V8 media for 60 hours. *A. castellanii* are added to the center of the lawn. *A. castellanii* consume the cells across the plate for a period of 12 or 18 days (based on amoeba activity). *Cryptococcus* strains able to resist amoeba have smaller areas of consumption when imaged. **B.***A. castellanii* in co-culture with *C. neoformans* expressing RFP on solid V8 media. White arrows indicate examples of *C. neoformans* - amoeba interaction. **C.** Boxplots representing amoeba resistance phenotypes for a diverse set of *C. neoformans* strains after 18 days of amoeba co-culture. The x-axis displays the strains assayed. The y-axis represents the area the amoeba consumed. Smaller clearance areas indicate greater resistance to amoeba.

This amoeba resistance assay was applied to a diverse set of *C. neoformans* strains that represent major sub-lineages within this species (Table 1). This assay revealed extensive variation in amoeba resistance between strain backgrounds (Fig 1C). Notably, there is no simple relationship between the site of collection and amoeba resistance; clinical strains isolated from patient samples exhibited both resistant and sensitive amoeba resistance phenotypes.

**Table 1.** Genetically diverse *C. neoformans* strains surveyed for amoeba resistance.

| Strain | Lineage | Mating Type | Site | Source |
| --- | --- | --- | --- | --- |
| Bt102 | VNBI | $\alpha$ | clinical | Litvintseva et al. (2003) |
| Bt22 | VNBI | $\alpha$ | clinical | Litvintseva et al. (2003) |
| Bt45 | VNBI | **a** | clinical | Litvintseva et al. (2003) |
| Ftc192-1 | VNBI | **a** | environmental | Chen et al. (2015) |
| Ftc267-1 | VNBI | $\alpha$ | environmental | Chen et al. (2015) |
| FTc555-1 | VNBI | **a** | environmental | Chen et al. (2015) |
| Bt1 | VNBII | $\alpha$ | clinical | Litvintseva et al. (2003) |
| Bt103 | VNBII | $\alpha$ | clinical | Litvintseva et al. (2003) |
| Bt206 | VNBII | **a** | clinical | Litvintseva et al. (2003) |
| Bt65 | VNBII | **a** | clinical | Litvintseva et al. (2003) |
| Bt75 | VNBII | $\alpha$ | clinical | Litvintseva et al. (2003) |
| AD1-7a | VNI | $\alpha$ | clinical | Dromer et al. (2007) |
| Bt130 | VNI | **a** | clinical | Litvintseva et al. (2003) |
| H99 | VNI | $\alpha$ | clinical | Perfect et al. (1980) |
| KN99a | VNI | **a** | laboratory | Nielsen et al. (2003) |

### Mapping Populations and Genome Sequencing

From our collection of genetically diverse strains, strains of opposite mating type (*MAT* **a**-*MAT α*) were identified that differed in their resistance to amoeba. We carried out pairwise mating tests to identify strain pairs with sporulation and germination efficiency suitable for establishing a large genetic mapping population. *C. neoformans* strains Bt22 and Ftc555-1 were chosen for further analysis based on spore viability and differences in their amoeba resistance. The low-resistance strain Bt22 (*MAT* **a**) is a clinical isolate while the high-resistance strain Ftc555-1 (*MAT α*) is an environmental isolate collected from a mopane tree; both strains were collected in Botswana [46]. By manual spore dissection, we isolated 384 progeny from a cross between these two strains. The genomes of these progeny were then sequenced on the Illumina NovaSeq 6000 platform to an average depth of ∼15*×*. Based on the resulting sequence data, the progeny were filtered based on criteria including sequencing depth, read quality, elevated ploidy, and clonality. After filtering, the final mapping population was composed of 304 recombinant progeny. 46,670 variable sites were identified between the parental strains that were collapsed into 4,943 haploblocks.

### Cross-Species Amoeba Resistance QTL

The F_1_ segregants generated from the Bt22 *×* Ftc555-1 cross exhibited a diverse response to amoeba predation (mean of predation area 23.46 cm^2^ and SD 14.99 cm^2^). There is a substantial amount of transgressive segregation – 16.9% of the segregants exhibited resistance higher than Ftc555-1, and 38.4% of segregants exhibited lower resistance than Bt22 (Fig 2B). Substantial transgressive segregation suggests that epistatic interactions between parental alleles contribute to both increased and decreased resistance beyond the parental phenotypes. Segregant genotypes and phenotypes were combined to carry out QTL mapping based on a marker regression approach [47]. This QTL analysis revealed that genetic variation for amoeba resistance in the mapping population is dominated by a single, major effect locus on chromosome 8 (Fig 3A). Segregants with Bt22 haplotypes at the chromosome 8 QTL peak exhibited significantly larger zones of amoeba clearance than those offspring with the Ftc555-1 haplotype (Fig 3B). The QTL on chromosome 8 explains an astonishing 62% of variation in amoeba resistance.

**Fig 2.**
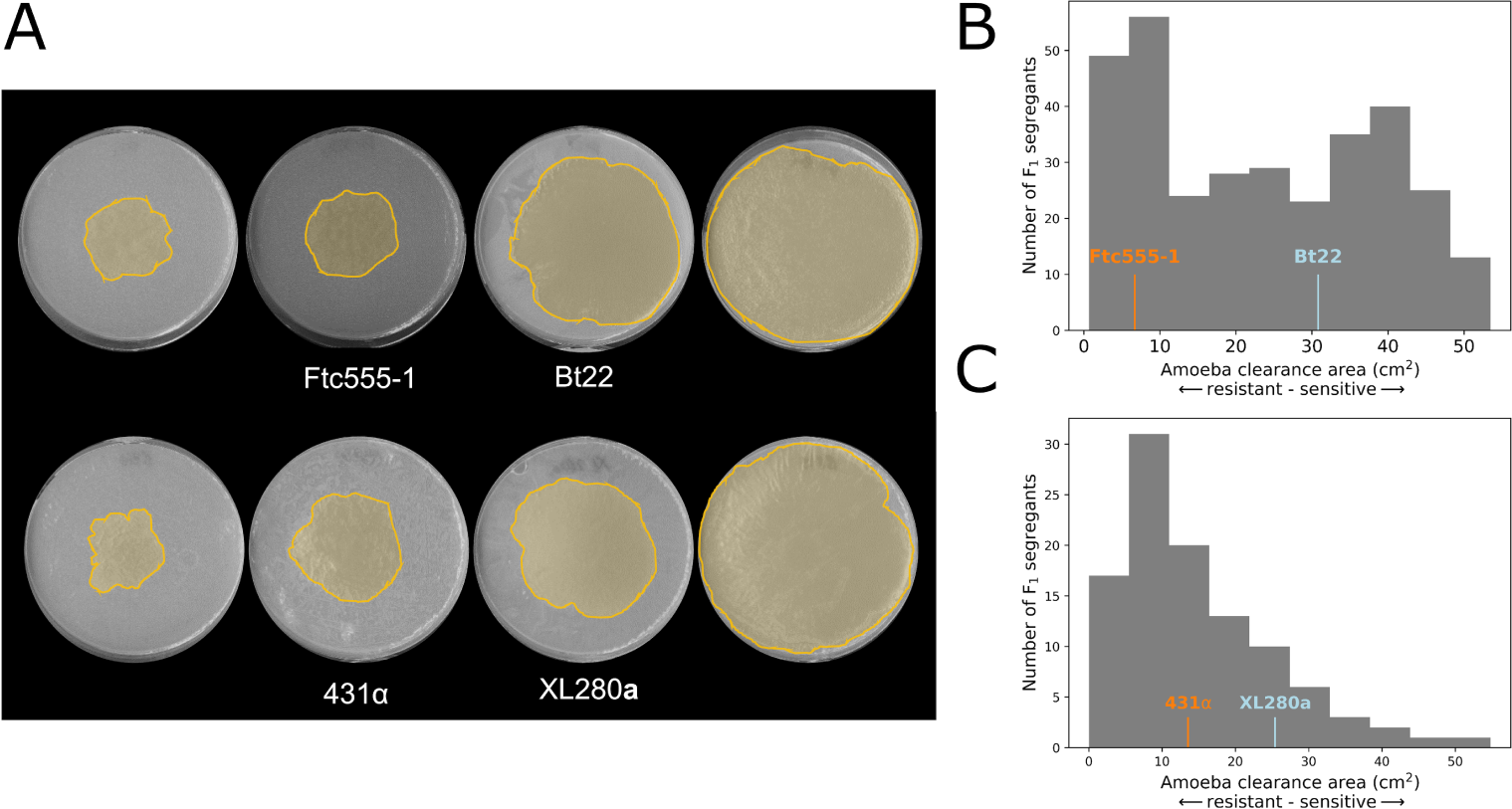
Phenotypic variation in amoeba resistance in mapping populations derived from both *C. neoformans* and *C. deneoformans* crosses. **A.** Representative images of plates from amoeba resistance assays. On the plates, the area consumed by amoeba are highlighted in yellow. Parental strains are shown in the middle panels, and transgressive segregants are on the left and right. **B.** A histogram displaying amoeba resistance of segregants in the *C. neoformans* cross. The x-axis represents the amoeba clearance area. Phenotypes of the two parental strains are indicated in orange (Ftc555-1) and blue (Bt22). **C.** A histogram of the *C. deneoformans* F_1_ progeny amoeba phenotypes. Phenotypes of the two parental strains are indicated in orange (431) and blue (XL280).

**Fig 3.**
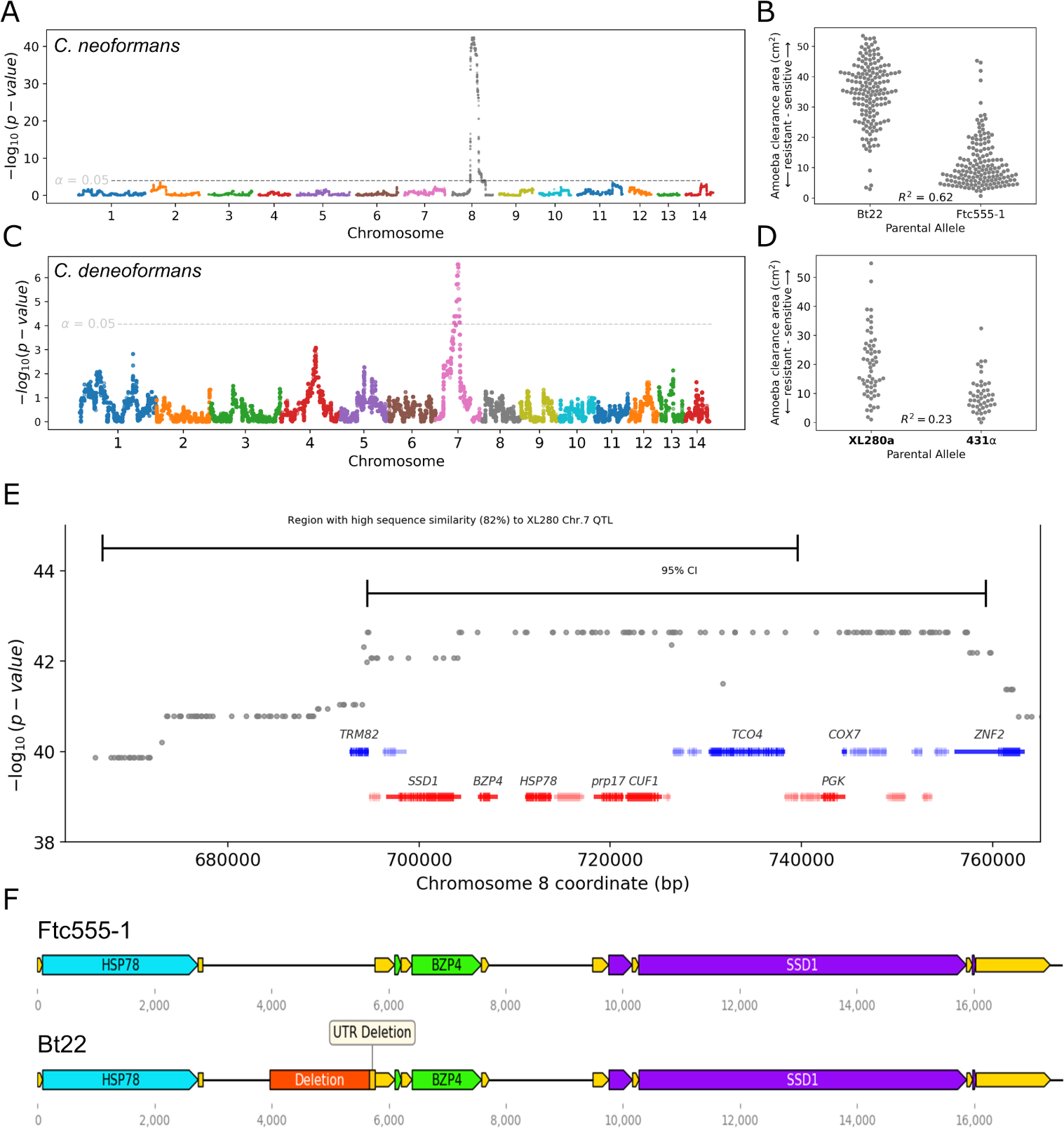
Amoeba resistance QTL for *C. neoformans* and *C. deneoformans*. **A.** Manhattan plot representing the association between genotype and amoeba resistance in the *C. neoformans* mapping populations. The dotted line indicates the significance threshold by permutation. **B.** Distributions of segregant phenotypes associated with the QTL peak on chromosome 8 for *C. neoformans*. The x-axis represents allelic state at the QTL peak. This peak explains 62% of the amoeba resistance variation (R^2^). **C.** Manhattan plot for amoeba resistance in the *C. deneoformans* mapping population. This peak explains 62% of the amoeba resistance variation (R^2^). **D.** Segregant phenotypes by chromosome 7 genotype for *C. deneoformans*. **E.** A magnified view of the 95% confidence interval of the *C. neoformans* QTL for amoeba resistance. Barred lines at the top are the 95% confidence intervals for the *C. deneoformans* and *C. neoformans* amoeba resistance QTL. **F.** Gene diagrams for the region around the *BZP4* gene for Ftc555-1 and Bt22. UTRs are shown in yellow.

To determine if there are similarities in the genetic architecture of amoeba resistance between closely related pathogenic species of *Cryptococcus*, we carried out a similar analysis using a mapping population derived from a *C. deneoformans* cross described in [47]. This cross, between strains XL280**a** and 431α, consists of 90 recombinant progeny. XL280**a** and 431α have only modest differences in amoeba resistance, but similar to the findings in *C. neoformans*, the *C. deneoformans* offspring exhibited a high degree of transgressive segregation for this trait. 15.4% of offspring displayed negative transgressive segregation (segregants with lower amoeba resistance than XL280**a**), 54.8% positive transgressive segregation (segregants with lower amoeba resistance than 431α), and 29.8% non-transgressive (Fig 2A,C). QTL analysis of this population identified a significant peak on chromosome 7 that explains 23% of the variance of amoeba resistance (Fig 3C). Progeny with the 431α allele on chromosome 7 have a higher average resistance to amoeba (Fig 3D). By examination of the genes under the QTL peak on chromosome 7, this region was found to be orthologous to the *C. neoformans* QTL peak on chromosome 8. These two regions share 82% nucleotide sequence identity and conserved synteny (Fig 3E), suggesting there are conserved genes required for amoeba resistance within the *Cryptococcus* species complex.

### Identification of an Amoeba Resistance Gene

To identify candidate causal variants for the QTL region on chromosome 8 in the *C. neoformans* cross, we analyzed the predicted effect of nucleotide sequence differences on annotated features in this region (S1 Table). 25 genes were within the identified 64 kb region that comprises the 95% confidence interval for this QTL. Ten of these genes have an annotated function or have homology to annotated genes in other fungal species. Across all of the genes in the QTL region, 31 synonymous mutations, 29 non-synonymous mutations, and two indels were identified. The two indels both result in nonsense mutations but the predicted genes in which they occur have no characterized function and no known gene deletion phenotype. Characterization of genetic variation in non-coding regions in the QTL region led to the identification of a large sequence difference between Bt22 and Ftc555-1, a 1789 bp deletion occurs in the Bt22 background in the intergenic region between the *BZP4* and *HSP78* genes. The Bt22 variant truncates 100 bp of the annotated *BZP4* 5’ UTR and 1689 bp further upstream (Fig 3F).

With *C. neoformans* phylogenetic data and short-read sequence data from [48], a strain was identified, Bt45, that is nearly genetically identical to Bt22 (∼200 SNP differences) but does not share the deletion upstream of *BZP4* (Fig 4A). Like Bt22, Bt45 is a clinical isolates collected in Botswana from a patient with HIV/AIDS in the early 2000s [44, 49]. Bt45 was found to exhibit significantly greater amoeba resistance than Bt22 (pairwise t-test, p < 0.0005), though not to the level observed for Ftc555-1 (Fig 4B). This comparison between these two nearly genetically identical strains is analogous to an “allele exchange” experiment and provides strong evidence that the non-coding variant identified upstream of *BZP4* is the likely causal variant underlying the Chromosome 8 QTL.

**Fig 4.**
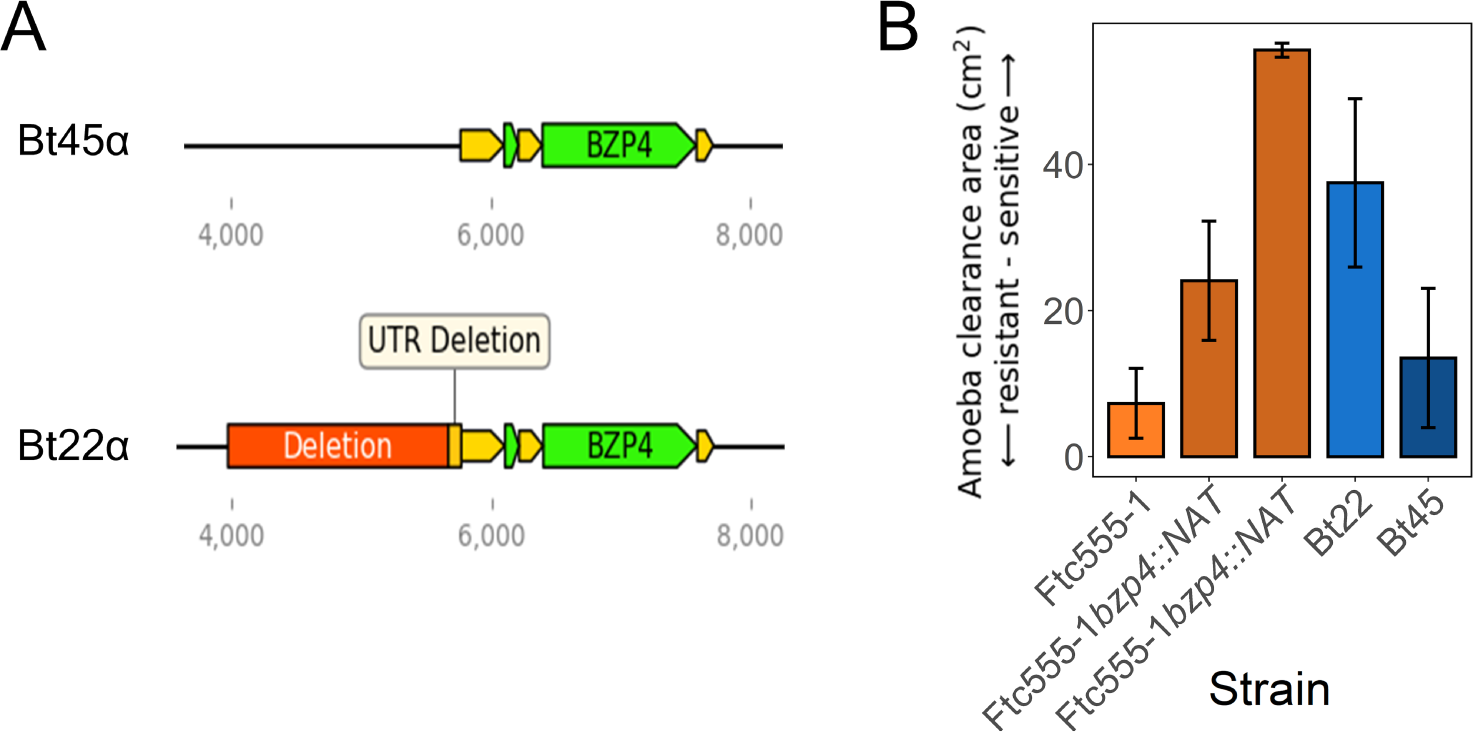
Disruption of *BZP4* reduces amoeba resistance. **A.** Models of the genomic regions surrounding *BZP4* in Bt22 and Bt45. **B.** Amoeba resistance assay for two independent gene deletions of *BZP4* in the Ftc555-1 background and a closely related strain of Bt22, Bt45. Bars are colored for comparison between closely related strains. The x-axis represents the strain tested. The y-axis represents the area amoeba consumed.

To provide further evidence of *BZP4*’s contribution to amoeba resistance, CRISPR-Cas9 editing was utilized to delete *BZP4* in the Ftc555-1 background [50, 51]. Two independent *bzp4*Δ mutants were isolated and their amoeba resistance phenotypes were assessed. Both mutants exhibited a significant reduction in amoeba resistance (pairwise t-test, p < 0.0005) and melanization (Fig 4B, S3 Fig).

In sum, multiple lines of evidence suggest that the 1789 bp deletion we identified upstream of *BZP4* is the causal variant for the large effect amoeba resistance QTL we identified on chromosome 8. For the sake of conciseness, in the text that follows the two allelic states at this locus are referred to as *BZP4^B^* (Bt22 allele) and *BZP4^F^* (Ftc555-1 allele).

### BZP4 is a pleiotropic QTG for Amoeba Resistance and Melanization

*BZP4* is a transcription factor that has been shown to play a role in regulation of the melanin synthesis pathway under nutrient deprivation conditions [52]. Variation at the *BZP4* locus was previously identified in a genome-wide association study (GWAS) for melanization [48]. In that study, *bzp4* loss-of-function mutations, found exclusively in clinical isolates, were shown to correlate with reduced melanization. A later study found that decreased expression of *BZP4* is correlated with decreased melanization in VNI clinical isolates [53].

Based on the role of *BZP4* in the regulation of melanin synthesis, we reasoned that the *BZP4* variant identified in the amoeba resistance mapping might also result in differences in the ability to produce melanin. While neither of the parent strains in the cross lacks melanin, Bt22 exhibits less melanin pigmentation than Ftc555-1 when grown under the same inducting conditions (Fig 5A). The *C. neoformans* mapping population was assayed for the ability to produce melanin when grown on L-DOPA plates. Segregants in the cross ranged from completely white (devoid of melanin) to a deep ebony color accompanied by melanin leaking into the surrounding media (Fig 5A). Across the segragants, 23.45% of the segregants displayed positive transgressive segregation (more melanized than Ftc555-1) while 24.43% displayed negative transgressive segregation (less melanized than Bt22). When the joint distribution of amoeba resistance and melaninization phenotypes among the offspring was assessed, a positive but non-linear relationship was observed (Fig 5D).

**Fig 5.**
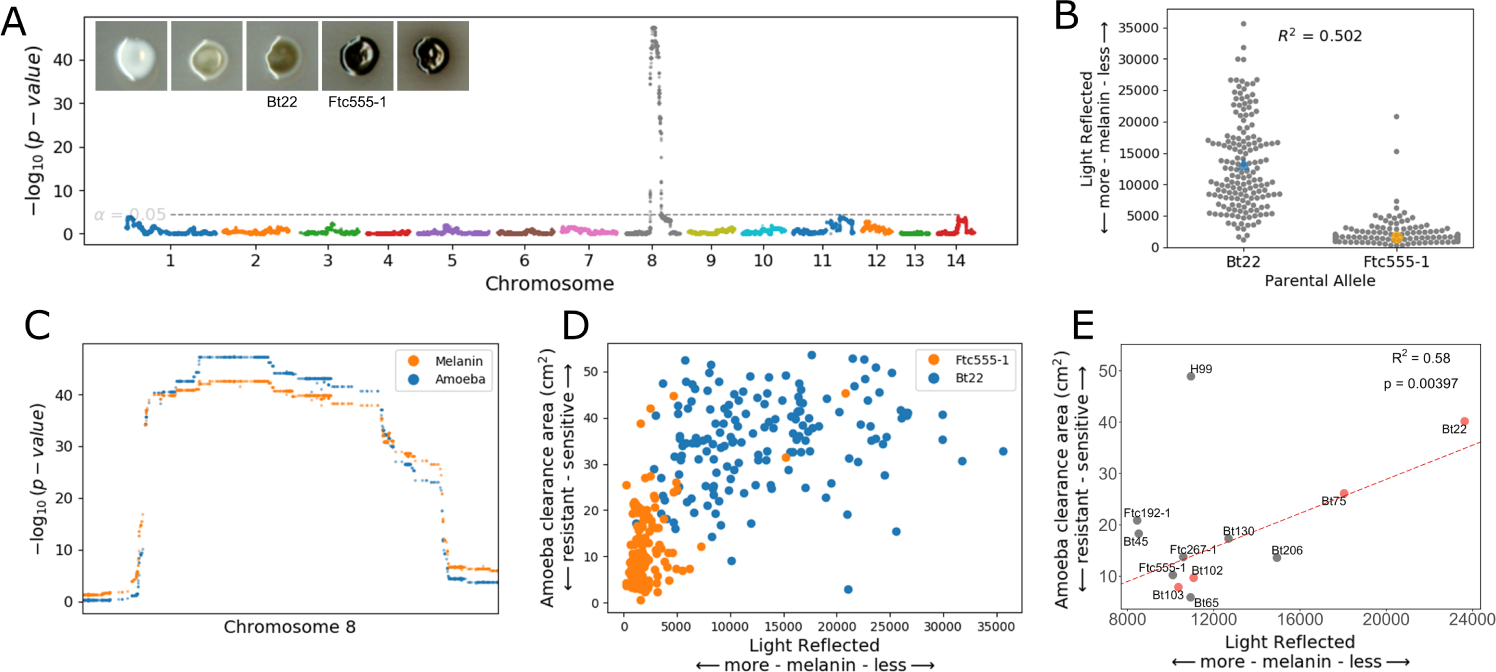
Melanization and amoeba resistance share the same QTL. **A.** Manhattan plot representing the association between genotype and melanization in the *C. neoformans* plot. The y-axis represents the strength of the association between genotype and light reflected (the degree of melanization). The x-axis represents the genomic location of the haploblocks used in the associations. The dotted line represents a significance threshold determined by a permutation test. Representative images of parents and segregants on L-DOPA media are also included. Each colony is from the randomized plates employed for QTL mapping. Images are brightened 30% to better display the difference in pigmentation. **B.** Segregant phenotypes at the maximum significance value of the QTL on chromosome 8. The x-axis represents the segregant allele at the maximum significance of the QTL. The y-axis represents the light reflected off of the colony when melanized. Parental strains are indicated in orange (Ftc555-1) and blue (Bt22). **C.** A magnified view of the chromosome 8 QTL peaks for amoeba resistance and melanization illustrating the QTL overlap. **D.** Plot comparing amoeba resistance and melanization phenotypes. The x-axis represents melanization and the y-axis represents amoeba resistance. Each dot represents a single segregant. Segregants are colored by their allele at the chromosome 8 QTL. **E.** Relationship between amoeba resistance and melanization in natural isolates. The linear regression line, and the coefficient of determination, are indicated for the linear model fit to the data with the strain H99 (outlier, top left) excluded.

QTL mapping based on the melanization phenotypes identified a major peak on chromosome 8 nearly identical in location to the QTL for amoeba resistance (Fig 5C). This QTL explains a remarkable 50.2% of the phenotypic variation for melanization (Fig 5B). Based on the similarity of the QTL for amoeba resistance and melanization (Fig 5C), as well as the previously demonstrated role of *BZP4* in the regulation of melanin synthesis, we propose that the non-coding deletion upstream of *BZP4* has pleiotropic effects on both of these traits.

In contrast to the findings in *C. neoformans*, the chromosome 7 QTL for amoeba resistance in the *C. deneoformans* cross does not appear to have a pleiotropic effect on melanization. Instead, a nonsense mutation in the gene *RIC8* is primarily responsible for variation in melanization for this cross as described in an earlier study from our research groups [47].

To test whether the relationship observed between amoeba resistance and melanization in our *C. neoformans* mapping population holds more broadly, we again employed the genetically diverse collection of *C. neoformans* strains described above. This collection was supplemented with additional strains that have predicted *BZP4* loss-of-function mutations, and amoeba resistance and melanization for each isolate was measured. All strains, including those with predicted *BZP4* loss-of-function mutations, are capable of producing melanin given sufficient incubation time (S3 Fig). However, large differences in the rate of melanization between strains were observed and images taken at two days of growth as a measure of the variation were used.

When comparing amoeba resistance and melanization across the diverse strain set, the reference strain H99 is notable as an outlier in terms of the bivariate relationship among these traits (S4 Fig). With H99 excluded, there is a strong linear relationship between amoeba resistance and melanization (R^2^ = 0.58) (Fig 5E). A notable trend among strains with predicted *BZP4* loss-of-function mutations is that those that melanize more readily (Bt103 and Bt102) are more resistant to amoeba than those that melanize slowly (Bt22 and Bt75) (Fig 5E).

### Gene Expression Differences Associated with *BZP4* Allelic Variation

Because the *BZP4* allele identified involves a deletion of a large upstream non-coding region, we hypothesized that the phenotypic effects of this allele are mediated by a reduction in the expression of the *BZP4* gene, with consequent effects on the downstream targets of this transcription factor. To test this hypothesis, gene expression was profiled with RNAseq. Using six offspring with each *BZP4* genotype (6 for Bt22 and 6 for Ftc555-1; 12 strains in total) transcriptional responses when grown on V8 medium were compared with or without the addition of amoeba.

*BZP4* was significantly differentially expressed between strains with the *BZP4^B^* and BZP4*^F^* alleles, in both amoeba and non-amoeba conditions (Fig 6A, B). *BZP4^B^* strains exhibited an average 1.83-Log_2_ fold decrease in expression relative to *BZP4^F^* strains when co-cultured with amoeba and a 2.06-Log_2_ fold decrease when amoeba were absent (Fig 6A). No other gene within the chromosome 8 QTL showed statistically significant differences in expression. While there are differences in *BZP4* expression between genotypes, no significant change in the expression of *BZP4* was observed between control and amoeba conditions (Fig 6A). This suggests that the effect of the *BZP4* allelic differences identified is not specific to the amoeba challenge conditions of our assay.

**Fig 6.**
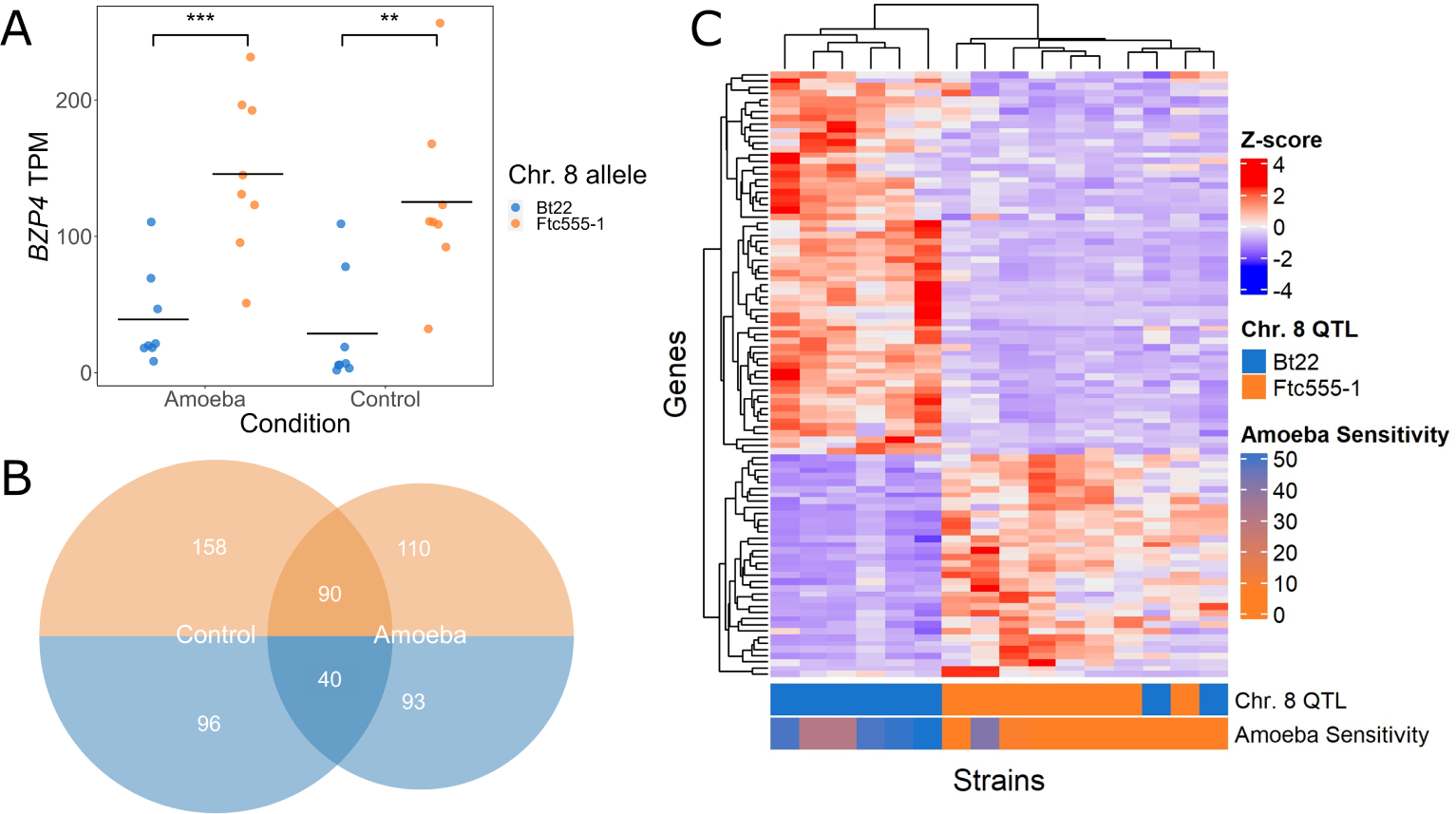
*BZP4* expression differs significantly between genotypes. **A.** Points representing the difference in transcript abundance between conditions in transcript per million (TPM) in segregants with each parental allele under the chromosome 8 QTL. The y-axis represents the total transcript counts. Boxplots are colored by parental allele. Significance is determined by ANOVA. “**” p < 0.005; “***” p < 0.0005. **B.** A Venn Diagram displaying the number of genes with increased and decreased expression for amoeba and control conditions based on the parental allele. Genes with decreased expression are colored in orange and those with increased expression are colored in blue. **C.** A hierarchically clustered heatmap of genes that strongly correlate with *BZP4* expression and are differentially expressed based on the *BZP4* allele in amoeba treated samples. Rows represent genes. Columns represent individual strains (parents and segregants). The colors of each cell represents z-scores. Colored rows at the bottom represent strains by their *BZP4* genotype and their amoeba sensitivity (area of clearance, measured in cm^2^).

Given that *BZP4* was identified as a candidate QTG for melanization, and the transcription factor it encodes has been previously implicated in the regulation of melanin synthesis genes, we predicted that such genes would also exhibit differences in expression as a function of *BZP4* genotype. Contrary to this prediction, we found that no major melanin synthesis genes were significantly differentially expressed between genotypes (when filtered for biological significance by fold change), when measured on the V8 growth media used in the amoeba experiments (Table S2).

Investigating genome wide expression differences, 587 genes exhibited a greater than 2-fold difference between genotypes in either amoeba and control conditions. Of the 587 differentially expressed genes, 130 are shared between conditions, 254 are specific to the control conditions, and 203 are amoeba specific (Fig 6B). Using GO term analysis for the 90 shared genes that have increased expression when *BZP4* expression is reduced, we find transmembrane transporter activity, oxidoreductase activity, and transition metal ion binding to increase in activity with reduced *BZP4* expression. Interestingly, there is a paucity of GO predictions for the 40 shared genes that are decreased in expression with reduced *BZP4* expression, as many of the genes are hypothetical or poorly characterized. The number of genes that have an inverse transcriptional relationship with *BZP4* expression indicates a role in gene repression. GO terms specific to amoeba conditions further include transmembrane transporter activity and oxidoreductase activity.

To identify genes potentially regulated by Bzp4 itself, we further focused on genes whose expression is strongly (anti-)correlated with *BZP4* (Pearson correlation, *|R| ≥* 0.5) and that show differential expression between strains with the *BZP4^B^* and *BZP4^F^* alleles (log_2_-fold change > 1 and *p_adj_ ≤* 0.05). This subsetting identified 36 genes that are positively correlated with *BZP4* and 62 genes that are negatively correlated (Fig 6C, Table S3). Hierarchical clustering of these genes, illustrated in Figure 6C, emphasizes the highly distinct expression patterns that these genes exhibit between strains with high and low amoeba resistance. The GO profile of this gene set is highly similar to that of the broader set of differentially expressed genes.

To identify potential regulatory motifs in the promoter regions of genes with similar expression patterns to *BZP4*, we used the motif analysis tool XSTREME [54, 55] to analyze 1 kb regions upstream of the 98 *BZP4*-correlated genes (positive and negatively correlated genes analyzed separately). Using this approach we identified the enriched sequence motif CACAKGCWA (K = T/G, W = T/A; S1 Fig) which is found in 25% of the genes positively correlated with *BZP4*, including *BZP4* itself, compared to a background rate of <1% across sets of 500 random genes. This enriched motif bears similarity to the general E-box motif CANNTG typically bound by basic Helix-Loop-Helix (bHLH) transcription factors [56] and a particularly close match to the PBE-box motif, CACATG, which is bound by phytochrome interacting bHLH transcription factors [57]. No consistent sequence motif was identified among the negatively correlated genes.

### Population Genetics of *BZP4*

Providing a more detailed understanding of allelic variation at *BZP4* in *C. neoformans*, we reanalyzed *BZP4* sequence variation based on 387 sequenced *C. neoformans* strains, originally described in Desjardins et al. 2017 [48].

Desjardins *et al.* identified four strains (Bt75, Bt102, Bt103, Bt147) within the VNBI and VNBII lineages with *BZP4* loss-of-function (LOF) alleles. Our bioinformatics analysis similarly predicted LOF alleles for these four strains and identified three additional VNI strains with likely LOF alleles – Bt3, Bt107, and Bt156. All seven of these strains with predicted *BZP4* LOF alleles were isolated in clinical settings. Based on the phylogenetic relationships of these strains, and the specific locations of the stop-gain mutations, it is likely that each of these alleles arose independently.

To explore genetic variation at the *BZP4* locus more broadly, we calculated two widely used measures of nucleotide sequence variation: a) *π*, the average per-base number of variable sites between pairs of individuals in a population; and b) Tajima’s D, a summary statistic that can be used to identify genomic regions with an excess of rare alleles. Negative values of Tajima’s D, which indicates an excess of rare alleles, can be caused by selective sweeps but can also result from demographic phenomena such as population bottlenecks. We compared *BZP4* estimates of *π* and Tajima’s D to estimates of the same parameters for 177 other transcription factors [58] and to all other annotated genes on chromosome 8 on a per lineage basis (S4 Table). In the VNI lineage, rare *BZP4* alleles are somewhat elevated (Tajima’s D = −1.97) compared to other genes on chromosome 8 and somewhat elevated compared to other transcription factors (Table S4). There is no evidence for an overabundance of rare *BZP4* alleles in the VNBI or VNBII lineages compared to the chromosome 8 average or relative to other transcription factors.

For the VNBI lineage, there are sufficient numbers of environmental and clinical strains (74 and 112 strains respectively) to compare Tajima’s D by isolation source. Estimates of Tajima’s D for *BZP4* are lower in clinical strains (Tajima’s D = −1.47) compared to environmental strains (Tajima’s D = −0.66). However, the difference in Tajima’s D between these two groups is not unusual relative to other genes on chromosome 8 or compared to other transcription factors.

### Epistatic QTLs for Amoeba Resistance and Melanization

While the pleiotropic chromosome 8 QTL we identified in the *C. neoformans* cross explains a large portion of variation for both amoeba resistance and melanization, both traits show continuous rather than bimodal distributions and there is a large degree of transgressive segregation. These observations suggested that there are likely additional alleles, perhaps interacting epistatically with the major effect allele on chromosome 8, that contribute to phenotypic differences in both of these traits. To test for epistatic interactions, our mapping population was subdivided based on genotype at the chromosome 8 QTL peak, and we re-ran the QTL mapping procedure for each subpopulation (S2 Fig). For amoeba resistance, a single epistatic QTL was found on chromosome 5, exclusive to the segregants with the Ftc555-1a allele at the chromosome 8 QTL (S2 FigA). This epistatic chromosome 5 QTL explains 19% of the variation within that subgroup and it increases the overall variance explained for amoeba resistance to 64%. We identified two epistatic QTL for melanization, one in the segregants that have the Bt22 chromosome 8 QTL allele and the second in those that have the Ftc555-1 allele (Fig S2B). The epistatic QTL in the Bt22 background is found on chromosome 1 and explains 26% of the variation within that subgroup. The epistatic allele in the Ftc555-1 background occurs on chromosome 7 and accounts for a more modest 7.8% of variance. With these epistatic interactions included, the variance in melanization explained by all of the QTL identified increases to 56.4%. Evidence of epistasis for amoeba resistance and melanization highlights the importance of strain background and the impact of individual allelic differences on traits of interest.

### Comparing Amoeba Resistance and Virulence

The accidental pathogen hypothesis is based on the similarities between amoeba and macrophage interactions with *Cryptococcus*. Both amoeba and macrophages employ similar methods of detecting, phagocytosing, and degrading fungal cells but the question remains: does survival when challenged with one phagocyte relate to success with the other? To answer this question, we measured the intracellular proliferation rate (IPR) in macrophages of progeny from the Bt22 *×* Ftc555-1 cross. F_1_ progeny were chosen to represent opposite extremes in terms of their amoeba resistance phenotypes. These low and high resistance strains were assayed alongside the parental strains.

J774A.1 murine macrophages were infected with *C. neoformans* cells and the internal proliferation rate of yeast cells was measured using time lapse microscopy as described in the Methods. All F_1_ progeny assayed, regardless of *BZP4* genotypes and amoeba resistance phenotypes, showed similar macrophage internal proliferation rates (Fig 7A). Phagocytic index, another measure of yeast-macrophage interactions, also showed no association with *BZP4* allelic variation or amoeba resistance (S5 Fig). Thus, in contrast to the predictions of the accidental pathogen hypothesis, we do not observe a relationship between amoeba resistance alleles and survival in macrophages.

**Fig 7.**
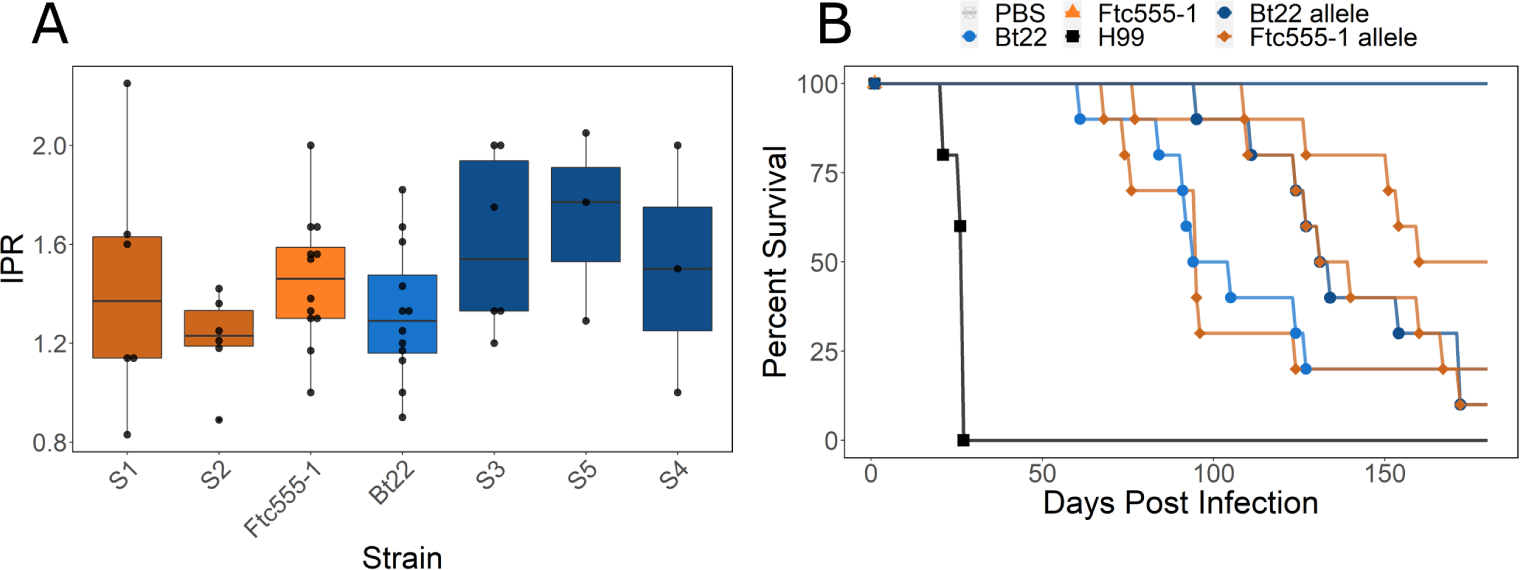
Amoeba resistance and measures of virulence are uncorrelated A. Barplots representing the internal proliferation rate of parental strains and segregants. Boxplots are colored by the chromosome 8 allele. Orange boxes have the Ftc555-1 allele and blue have the Bt22 allele, darker colors indicate segregants. Dots represent individual measurements. Strains are oriented in rank order of amoeba resistance (highest to lowest resistance). Significance determined by ANOVA (p = 0.29 F = 1.28). **B.** Survival curves for animals infected with the parental strains and a group of F_1_ segregants. H99 is in black, Ftc555-1 is in orange, and Bt22 is in blue. Segregant curves are colored by parental allele under the chromosome 8 QTL, dark blue represents the Bt22 allele and darker orange represents the Ftc555-1 allele.

To explore the relationship between amoeba resistance and the ability to cause disease in animal models, an equal number of 4-5 week old male and female A/J mice were intranasally infected with F_1_ progeny from the *C. neoformans* cross, the parental strains, and the reference strain H99. Survival was monitored for a period of 179 days, with animals sacrificed based on disease progression symptoms (Fig 7B). Though this analysis involve only a modest number of strains, we observed no relationship between *BZP4* genotype and virulence in mice. The virulence of the parental strains, Bt22 and Ftc555-1, is the opposite of their amoeba resistance phenotypes. Bt22, which exhibits low resistance to amoeba predation, has modest virulence with a time to 50% lethality (LT50) of ∼92 days. This is in stark contrast to Ftc555-1, which is highly resistant to amoeba, but is completely avirulent in the mouse model of infection employed. The reference strain, H99, is strongly virulent in mice (LT50 ∼21 days) but has very low amoeba resistance. Furthermore, we detected no association between LT50 estimated from murine survival curves and the *BZP4* genotype of a small number of segregants (Fig 7B).

Finding a lack of correlation between amoeba resistance and virulence for strains from our mapping population, the analysis was broadened to include an additional nine, genotypically diverse *C. neoformans* strains. Using the same intranasal murine infection model described above, we found that virulence was highly variable among strains, but again found no correlation between amoeba resistance and LT50 measures of mouse survival (Fig S5) [59].

## Discussion

Our findings provide novel insights into the genetic architecture of fungal-amoebal interactions and the potential impact of selection for amoeba resistance on *Cryptococcus* virulence. Using QTL mapping, we identified a transcription factor, *BZP4*, that is important for *C. neoformans* survival in the presence of amoeba. This gene also affects melanization, a classical virulence trait that is considered important for the pathogenic abilities of *Cryptococcus* [32, 60–62]. Despite its role in mediating interactions with amoeba and the production of melanin, allelic variation at *BZP4* is not predictive of proliferation rates in macrophages or virulence in mouse models of infection. This suggests that the relationship between resistance to amoeba and virulence potential may be more complex than the accidental pathogen hypothesis predicts.

Our findings share a mix of both similarities and differences to a recent study by Fu *et al.* that employed experimental evolution to identify phenotypes and mutations selected for during *Cryptococcus* co-culture with amoeba [63]. A reduction in melanization was one of the phenotypic changes they observed that was most consistent across the three genetic backgrounds studied. This is in contrast to our findings, where we found that melanization was correlated with *higher* resistance to amoeba. However, similar to what we report here, their study failed to detect an association between amoeba resistance and macrophage challenge or virulence in mice. A particularly interesting genotypic change that Fu *et al.* identified in three independently evolved populations, derived from the H99 strain background, were duplications of chromosome 8. In light of our current study, the effect of a duplication of *BZP4*, which is located on chromosome 8, should be noted as a potential additional positive effect on amoeba resistance.

Studies by Idnurm and colleagues have also failed to find a relationship between amoeba resistance and virulence potential in Basidiomycete yeasts [41, 64, 65]. In these studies, experimentally evolving *C. neoformans* and *C. deneoformans* strains in the presence of *A. castellanii* led to the isolation of amoeba-resistant RAM mutants [41] Despite the increased resistance to amoeba, such RAM mutants show decreased virulence in mouse models of infection because the RAM pathway is required for viability at 37*^◦^*C [64]. Another recent study by the same group investigated the evolution of *Sporobolomyces primogenesis*, a non-pathogenic Basidiomycete yeast, in the presence of amoebae [65]. In this case, increased resistance was shown to be due to a loss-of-function mutation in the calcineurin pathway. Calcineurin signaling is considered to be essential for virulence in most pathogenic fungi, suggesting that genetic variation that favors escape from amoeba predation may, in some cases, decrease virulence potential in mammals.

A caveat that applies to our study as well as all prior investigations of the accidental pathogen hypothesis, are the challenges inherent to comparing virulence between *Cryptococcus* strains. Our analysis does not account for potentially important effects such as host genotype or experimental parameters such as the level of infective inoculum [66]. A second limitation comes from the use of a single species and strain of *Acanthamoeba* to assess resistance to amoeba. *Acanthamoeba* is commonly found in the same niches as *Cryptococcus* and is the primary organism used in studies of environmental predator-prey interactions with fungal pathogens [18, 19]; however, as a selective pressure, there are likely other amoeba that are present in these niches. A consideration of these and other complications point to the need to adopt a holistic approach as we attempt to define the ecological and evolutionary factors that select for virulence traits and increase pathogenic potential.

A striking outcome of our study is the discovery of an amoeba resistance QTL in homologous genomic regions for both *C. neoformans* and *C. deneoformans*. Cross species QTLs are rare, but they have been found for drought resistance between species of legume [67], a cardiovascular disease marker between humans and baboons [68], and gravitropism in corn and *Arabidopsis* [69]. Our study marks the first detection of cross species QTL in fungi and it suggests that the amoeba survival mechanisms discovered may be conserved between different pathogenic species of *Cryptococcus*. We have not, as yet, identified the specific causal variant for the amoeba QTL in *C. deneoformans*, though non-coding variants in the vicinity of *BZP4* are among the top candidates we intend to pursue in future work.

We were initially surprised to find a single QTL for amoeba resistance and melanization in *C. neoformans*. The continuous distribution of amoeba resistance and melanization in the F_1_ progeny implied more complex regulation than a single QTL would explain. Continuous traits are often governed by epistatic interactions between genes leading to the consideration of loci-loci interactions when discussing the effect of a QTL [70, 71]. Prior studies from our group have uncovered complex epistatic relationships that govern virulence phenotypes utilizing QTL mapping in *Cryptococcus* [47, 72]. Using the same techniques, we discovered an additional QTL for amoeba resistance and two additional QTL for melanization. Future work will detail the genes that underly these loci and their epistatic interactions. These additional epistatic QTL are indicative of the polygenic nature of stress response regulation. They also provide insight into the impact of strain background on the connection between individual loci and phenotype.

The importance of strain background prompted further investigation into the results of a prior GWAS analysis that implicated *BZP4* loss-of-function mutations with reduced melanization capacity [48]. In our re-analysis of the large set of sequenced strains from that study, we note that 9 out of 10 candidate *BZP4* loss-of-function alleles we identified come from strains isolated in clinical settings. In this regard, it is interesting to note that Yu et al. [53] found that reduced expression of *BZP4* is unique to the clinical strains they analyzed. Furthermore, each of the candidate *BZP4* loss-of-function mutations we identified appears to be both independent and recent based on comparison to closely related strains. These observations lead us to speculate that *BZP4* loss-of-function mutations may actually be advantageous during human infection which would further call into question the connection between amoeba resistance and virulence. However, one can not ignore the fact that *C. neoformans* strains represented in laboratory collections are biased towards clinical isolates; more intensive study of environmental isolates will be required to convincingly demonstrate variation that distinguishes pathogenic for non-pathogenic strains.

In our *C. neoformans* mapping population, we observed a positive, but non-linear relationship, between amoeba resistance and melanization, and *BZP4* is a candidate QTG for both of these traits. Melanization is understood to be important for macrophage resistance [73] primarily by increasing resistance to reactive oxygen and nitrogen species [74] and other stresses [31]. Furthermore, prior studies suggest that Bzp4 is a transcriptional activator of *LAC1* [52], the gene responsible for the enzyme laccase that catalyzes the reaction of dopamine to melanin in *Cryptococcus* [75] This raises the question, “Do differences in melanization mediate variation in amoeba resistance?” We found that *BZP4* is differentially expressed, as a function of *BZP4* genotype, in both control and amoeba conditions. However, *LAC1* expression, and the expression of other key genes in the melanin synthesis pathway, do not vary as a function of *BZP4* genotype in the V8 media conditions used to co-culture amoeba and *Cryptococcus*. In addition, we did not observe melanization of *Cryptococcus* on V8 media, regardless of the presence or absence of amoeba. This is in contrast to our finding that amoeba resistance broadly correlates with melanization, and the observation that *BZP4* loss-of-function strains that rapidly melanize have increased amoeba resistance. However, a lack of differential expression of *LAC1* does not rule out the possibility that laccase levels differ between the strains, and it is possible that melanin variation is manifested after phagocytosis and thus not readily observable in the V8 growth medium employed here. Deeper investigation into melanin regulation and synthesis during amoeba challenge will be necessary to tease apart the contributions of melanization to amoeba resistance.

Using RNA-seq transcriptional profiling we identified a core set of genes that are differentially expressed as a function of genotype at the *BZP4* locus and are strongly correlated/anti-correlated with *BZP4*’s expression. We hypothesize that such genes may be targets of Bzp4. Using motif analysis we identified a sequence motif that is enriched upstream of genes that are positively correlated with *BZP4*. The core of the identified sequence motif bears strong similarity to an E-box family motif found in plants called the PBE-box [57]. Although E-box motifs are typically bound by bHLH transcription factors, there is precedence for fungal bZip transcription factors binding to E-box like elements [76, 77]. Future studies that characterize the transcriptional targets of Bzp4, such as through ChIP-seq analysis, will be critical to identify genes regulated by Bzp4 and elucidate Bzp4’s role in regulating traits such as amoeba resistance, melanin synthesis, and capsule formation.

If the transcription factor *BZP4* is a key element in mediating escape from amoeba, as the evidence supports, an interesting question is whether *BZP4* function is an absolute requirement. Two clinical isolates, Bt102 and Bt103, suggest that this is not the case. These strains have predicted *bzp4* loss-of-function mutations, but were highly resistant to killing by amoeba. This could be due to allelic variation acting either downstream of *BZP4* or in a pathway parallel to *BZP4* that also governs amoeba resistance. For example, our analysis of epistasis identified a QTL region on chromosome 5 that further contributes to variation in amoeba resistance. The causal variants and corersponding genes underlying this locus have not yet been identified. In a similar vein, mapping studies that employ other genetic backgrounds are likely to identify additional variants and genes that contribute to amoeba resistance. It is a distinct possibility that variation at other such amoeba-relevant genes may be better predictors of mammalian virulence than Bzp4. This emphasizes the importance of strain background in understanding the broader implications of gene function. We note that a consideration of genetic background is important not only for studies of natural genetic variation but also for molecular genetic studies of gene function as the effects of even large scale perturbations such as gene deletions can vary both within and between closely related species [78–80].

In summary, the transcription factor Bzp4 is important for the survival of *Cryptococcus* when exposed to phagocytic amoebae. Despite the importance of *BZP4* in amoeba resistance, *BZP4* function is not correlated with survival in macrophages nor is it predictive of virulence in mice. Furthermore, *BZP4* function is frequently lost in clinical isolates of *Cryptococcus*. While interactions with amoebae cannot be ruled out as a contributing factor to the evolution of *Cryptococcus* virulence, our findings suggest that phagocytic amoebae and phagocytic immune cells, despite their many parallels, are distinct niches from the perspective of fungal survival.

## Supporting information

**S1 Fig. A DNA sequence motif enriched in upstream sequences of genes positively correlated with BZP4 expression.** The motif analysis tool XSTREME [54] was used to identify sequence motifs over-represented in 1 kb upstream regions of genes that exhibit expression similar to *BZP4*. This sequence logo represents a motif found in 9 of 36 upstream regions (E-value 1.56e-008).

**S2 Fig. Epistatic QTLs for amoeba resistance and melanization.** The offspring of the Bt22 *×* Ftc555-1 cross were subset by genotype at the chromosome 8 QTL, and the QTL mapping procedure was repeated for each subpopulation. Variation on chromosome 8 was excluded from consideration. The dotted lines indicate significance thresholds (*α* = 0.05) determined by permutation testing. **A.** Manhattan plots for QTL mapping of amoeba resistance, conditional on chromosome 8 QTL genotype. **B.** Manhattan plots for QTL mapping of melanization, conditional on chromosome 8 QTL genotype.

**S3 Fig. BZP4 mutants are slow to melanize.** Images of colonies pinned onto L-DOPA plates and imaged after two and three days of growth. The *bzp4* deletion mutant and Bt22, which has a 2 kb deletion upstream of *BZP4*, are slow to melanize but still capable of some degree of melanin synthesis. Images of colonies are uniformly brightened by 30% to better visually contrast the level of melanization.

**S4 Fig. Amoeba resistance does not predict virulence in mouse models of infection.** The relationship between amoeba resistance and median time to death (LT50) of mice infected with *C. neoformans* strains. Significance determined by linear regression. Segregants that were avirulent were assigned a value of 190 days for LT50.

**S5 Fig. *BZP4* genotypes do not predict phagocytic index in mammalian macrophages.** Boxplots representing the phagocytic index of parental strains and segregants. Strains are oriented in rank order of amoeba sensitivity. Boxplots are colored by chromosome 8 QTL genotype. Orange indicates strains with the Ftc555-1 allele and blue those with the Bt22 allele. Significance determined by ANOVA (F = 22.18; p<0.0001).

**S1 Table. SNPs under the chromosome 8 QTL peak.**

**S2 Table. Differential expression of melanin synthesis genes as a function of BZP4 genotype.** Differential expression of genes involved in the melanin synthesis pathway [52]. Mean expression and mean log2-fold change represent the difference in expression for strains with the Bt22 allele at the BZP4 locus relative to strains with the Ftc555-1 allele.

**S3 Table. Transcripts that are strongly correlated or anti-correlated with *BZP4* expression.** 98 genes (101 transcripts) were found to be differentially expressed and strongly correlated or anti-correlated (*|r| >* 0.5) with *BZP4* expression.

**S4 Table. Measures of sequence diversity for BZP4.** *π* and Tajima’s D for *BZP4*, for each of the three major lineages of *C. neoformans* compared to other genes on Chromosome 8 and other transcription factors (“control gene sets”). The fifth percentile of the distribution of Tajima’s D in the control gene sets is provided for comparison. The list of predicted transcription factors is from Jung et al. 2015 [58].

**S5 Table. Strains used in this study.**

**S5 Table README. Description of columns in S5 Table.** A plain text file describing the information in S5 Table.

## Supporting information

All supplementary figures

## Acknowledgements

We thank Dr. Calla Telzrow (Ph.D.), Dr. Corinna Probst (Ph.D.), and Dr. Andrew Alspaugh (M.D.), for their expert advice in RNAseq and metallo-ion biology in *Cryptococcus*. We also thank Dr. Arturo Casadevall (M.D., Ph.D.) for his comments and feedback on an earlier version of this manuscript.

Research reported in this publication was supported by the National Institute of Allergy and Infectious Diseases of the National Institutes of Health under award number R01AI133654. The content is solely the responsibility of the authors and does not necessarily represent the official views of the National Institutes of Health.

## Supplementary Figures

**Figure S1:**
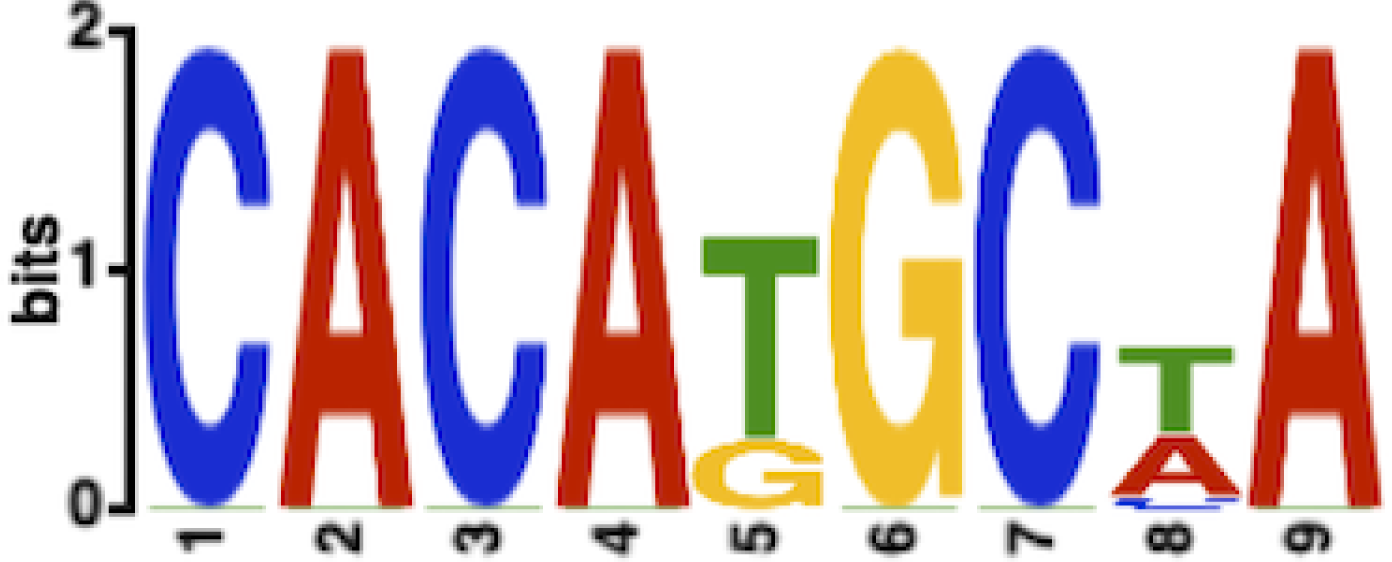
The motif analysis tool XSTREME (Grant and Bailey, 2021) was used to identify sequence motifs over-represented in 1 kb upstream regions of genes that exhibit expression similar to *BZP4*. This sequence logo represents a motif found in 9 of 36 upstream regions (E-value 1.56e-008)

**Figure S2:**
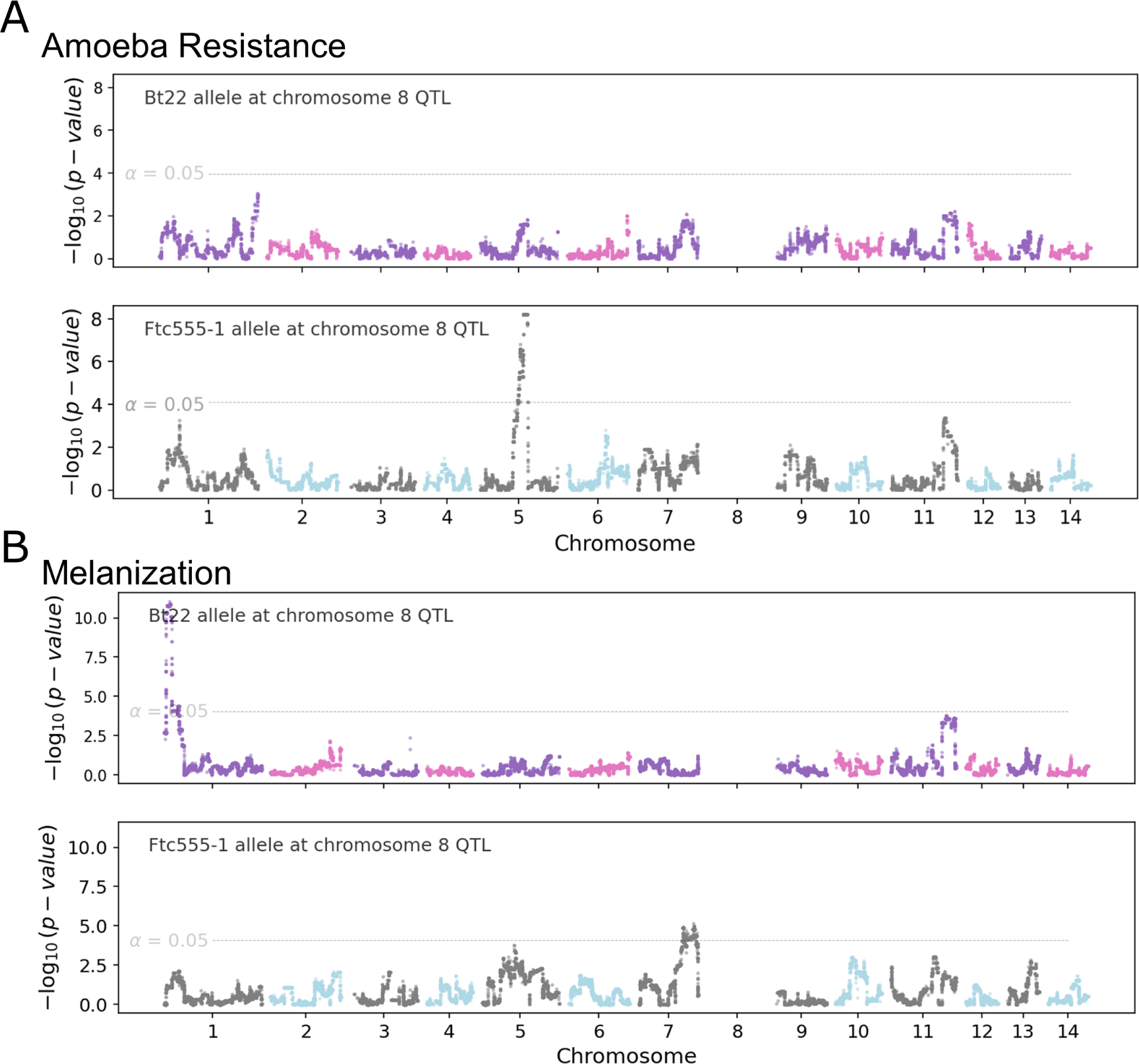
Epistatic QTLs for amoeba resistance and melanization. The offspring of the Bt22 *×* Ftc555-1 cross were subset by genotype at the chromosome 8 QTL, and QTL mapping was repeated for each subpopulation. Variation on chromosome 8 was excluded from consideration. The dotted lines indicate significance thresholds (*α* = 0.05) determined by permutation testing. **A.** Manhattan plots for QTL mapping of amoeba resistance, conditional on chromosome 8 QTL genotype. **B.** Manhattan plots for QTL mapping of melanization, conditional on chromosome 8 QTL genotype.

**Figure S3:**
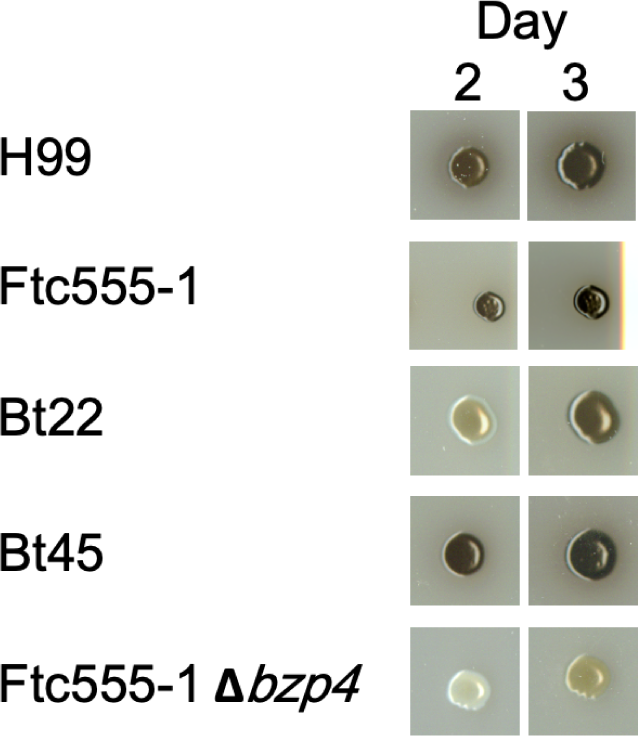
*BZP4* mutants are slow to melanize. Images of colonies pinned onto L-DOPA plates and imaged after two and three days of growth. The *bzp4* deletion mutant and Bt22, which has a 2 kb deletion upstream of *BZP4*, are slow to melanize but still capable of some degree of melanin synthesis. Images of colonies are uniformly brightened by 30% to better visually contrast the level of melanization.

**Figure S4:**
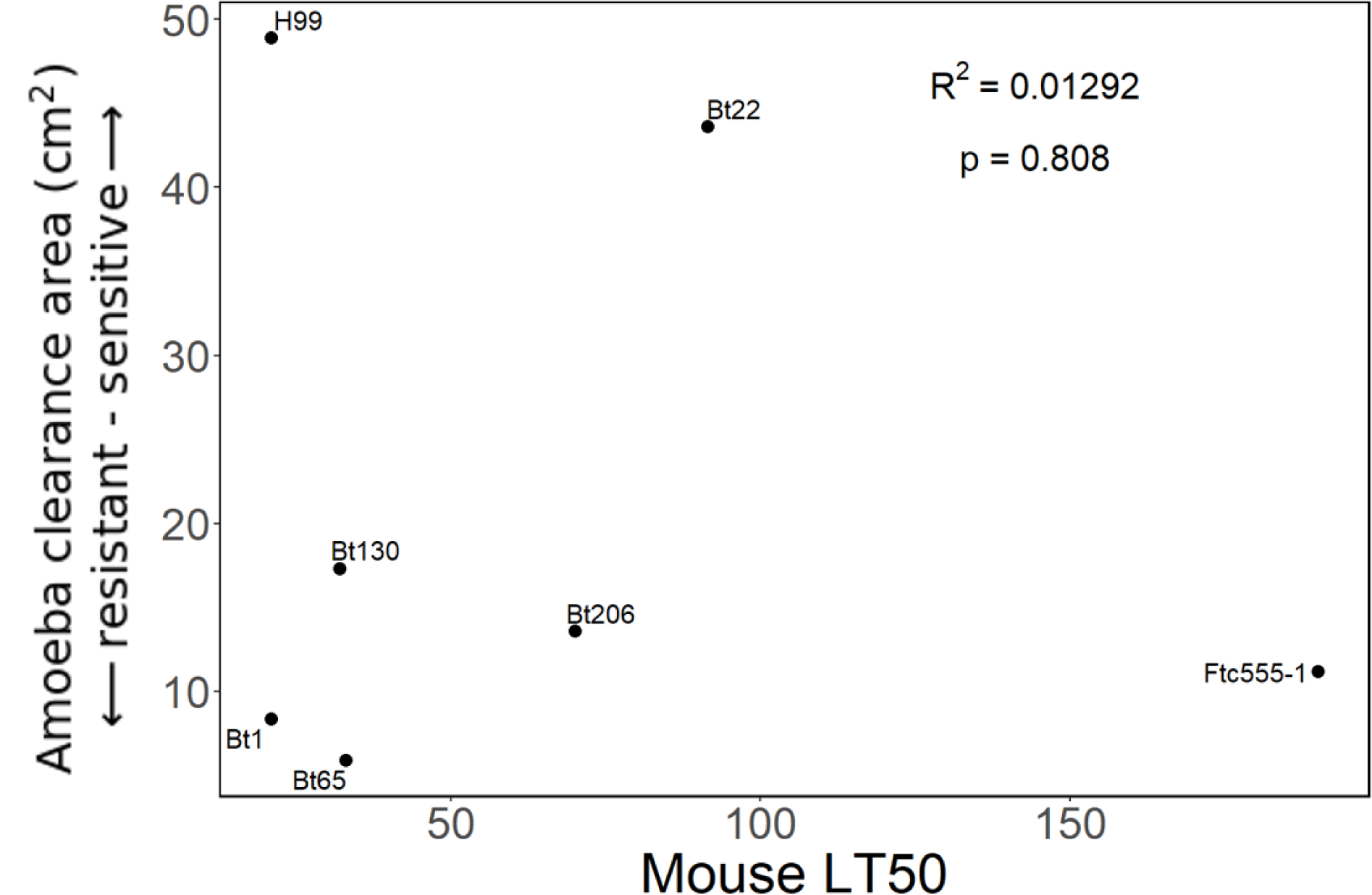
Amoeba resistance does not predict virulence in mouse models of infection. The relationship between amoeba resistance and median time to death (LT50) of mice infected with *C. neoformans* strains. Significance determined by linear regression. Segregants that were avirulent were assigned a value of 190 days for LT50.

**Figure S5:**
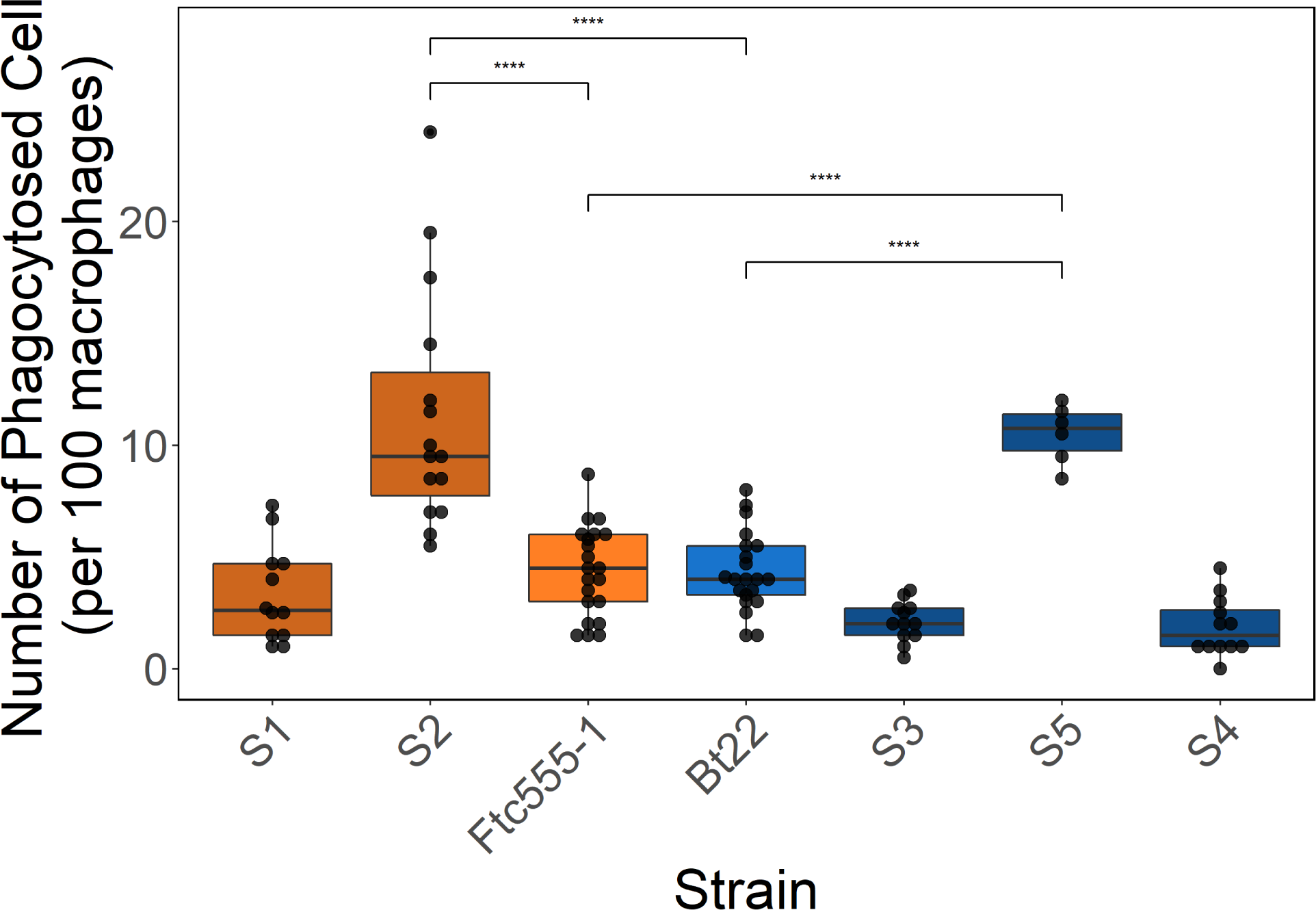
Phagocytic index is uncorrelated with parental genotype at the chromosome 8 QTL. Boxplots representing the phagocytic index of parental strains and segregants. Strains are oriented in rank order of amoeba sensitivity. Boxplots are colored by chromosome 8 QTL genotype. Orange indicates strains with the Ftc555-1 allele and blue those with the Bt22 allele. Significance determined by ANOVA (F = 22.18; p<0.0001).

## Notes

### Competing Interest Statement

The authors have declared no competing interest.

### Summary of Updates

Figure 6 is updated to include genes that correlate with BZP4 expression. The text on figure 6 has also been changed to incorporate the new results and better communicate that segregants were analyzed. Table 1 now includes mating type. Text has been added to the discussion to further clarify why we the study is unable to fully separate melanin and amoeba survival. A section on population genetics was added to further explore genetic variation at BZP4 using the metrics Tajima's D and pi.

