## Supplementary material for "Amoeba predation of *Cryptococcus*: A quantitative and population genomic evaluation of the Accidental Pathogen hypothesis": All supplementary figures

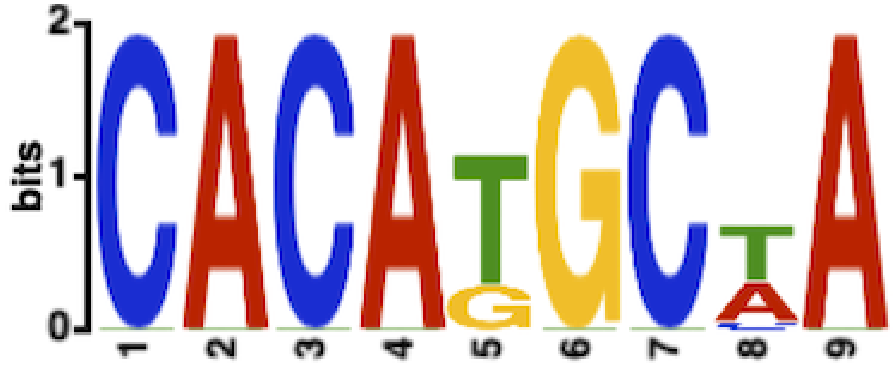

Figure S1: The motif analysis tool XSTREME (Grant and Bailey, 2021) was used to identify sequence motifs over-represented in 1 kb upstream regions of genes that exhibit expression similar to *BZP4*. This sequence logo represents a motif found in 9 of 36 upstream regions (E-value 1.56e-008)

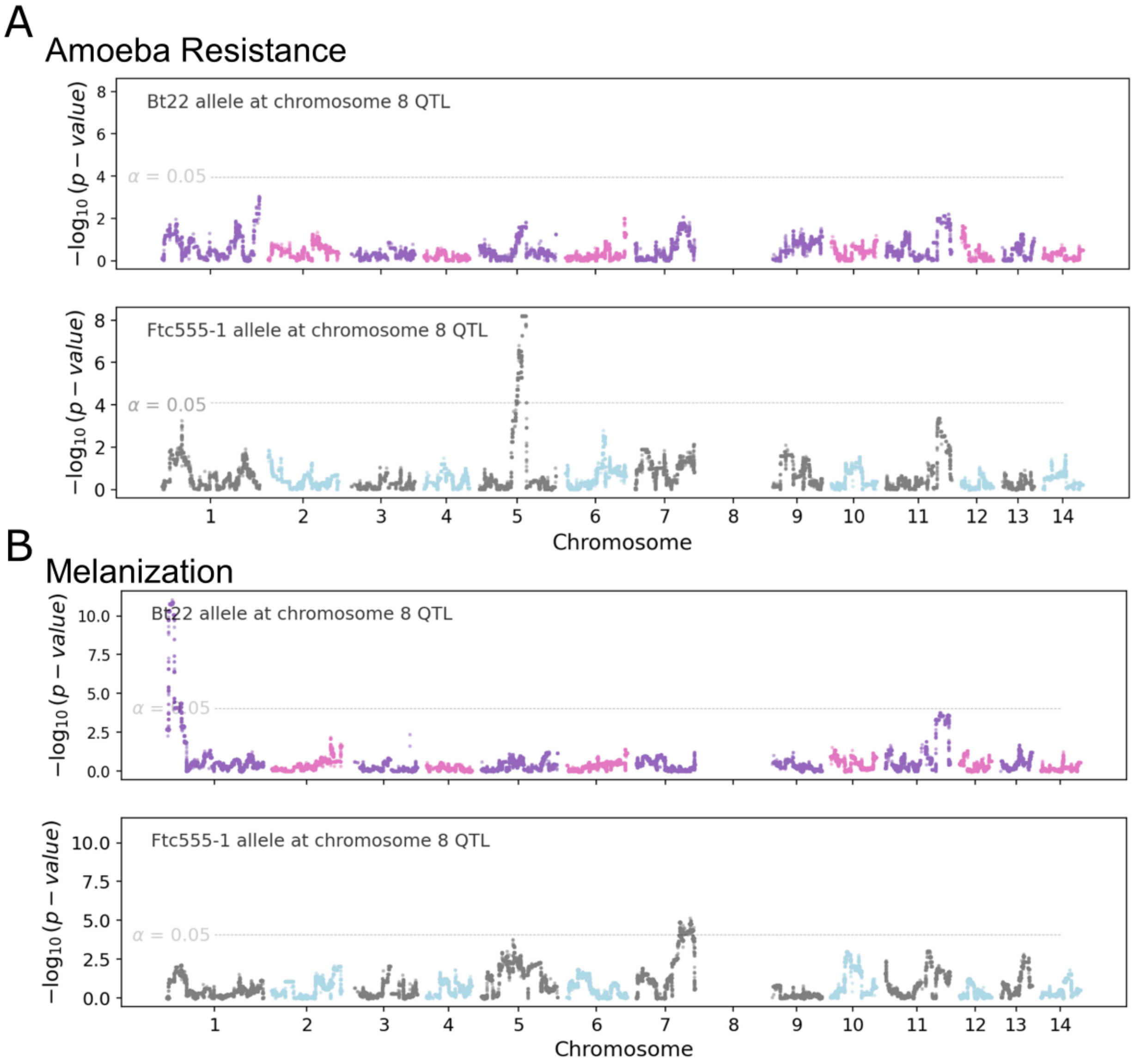

Figure S2: **Epistatic QTLs for amoeba resistance and melanization.** The offspring of the Bt22 $\times$ Ftc555-1 cross were subset by genotype at the chromosome 8 QTL, and QTL mapping was repeated for each subpopulation. Variation on chromosome 8 was excluded from consideration. The dotted lines indicate significance thresholds ($\alpha$ = 0.05) determined by permutation testing. **A.** Manhattan plots for QTL mapping of amoeba resistance, conditional on chromosome 8 QTL genotype. **B.** Manhattan plots for QTL mapping of melanization, conditional on chromosome 8 QTL genotype.

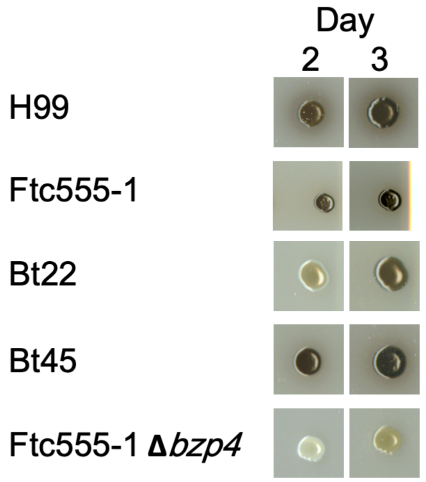

Figure S3: ***BZP4* mutants are slow to melanize** Images of colonies pinned onto L-DOPA plates and imaged after two and three days of growth. The *bzp4* deletion mutant and Bt22, which has a 2 kb deletion upstream of *BZP4*, are slow to melanize but still capable of some degree of melanin synthesis. Images of colonies are uniformly brightened by 30% to better visually contrast the level of melanization.

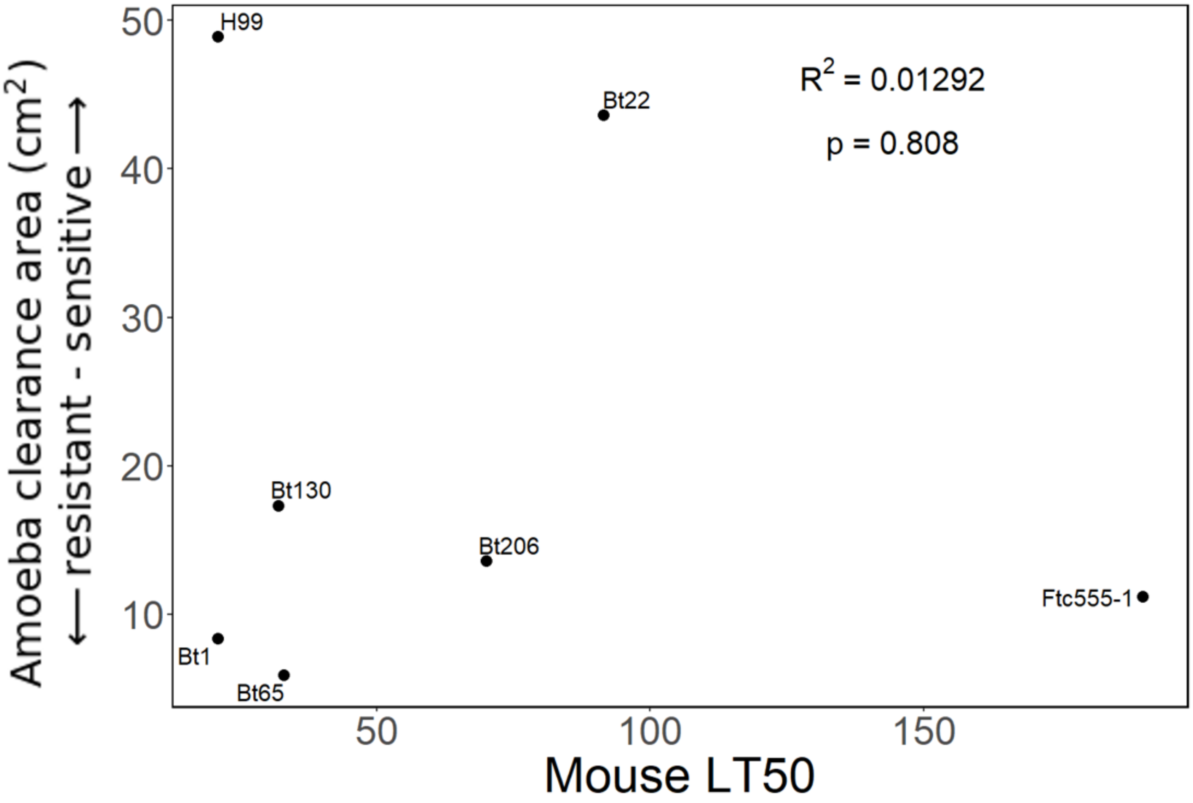

Figure S4: **Amoeba resistance does not predict virulence in mouse models of infection.** The relationship between amoeba resistance and median time to death (LT50) of mice infected with *C. neoformans* strains. Significance determined by linear regression. Segregants that were avirulent were assigned a value of 190 days for LT50.

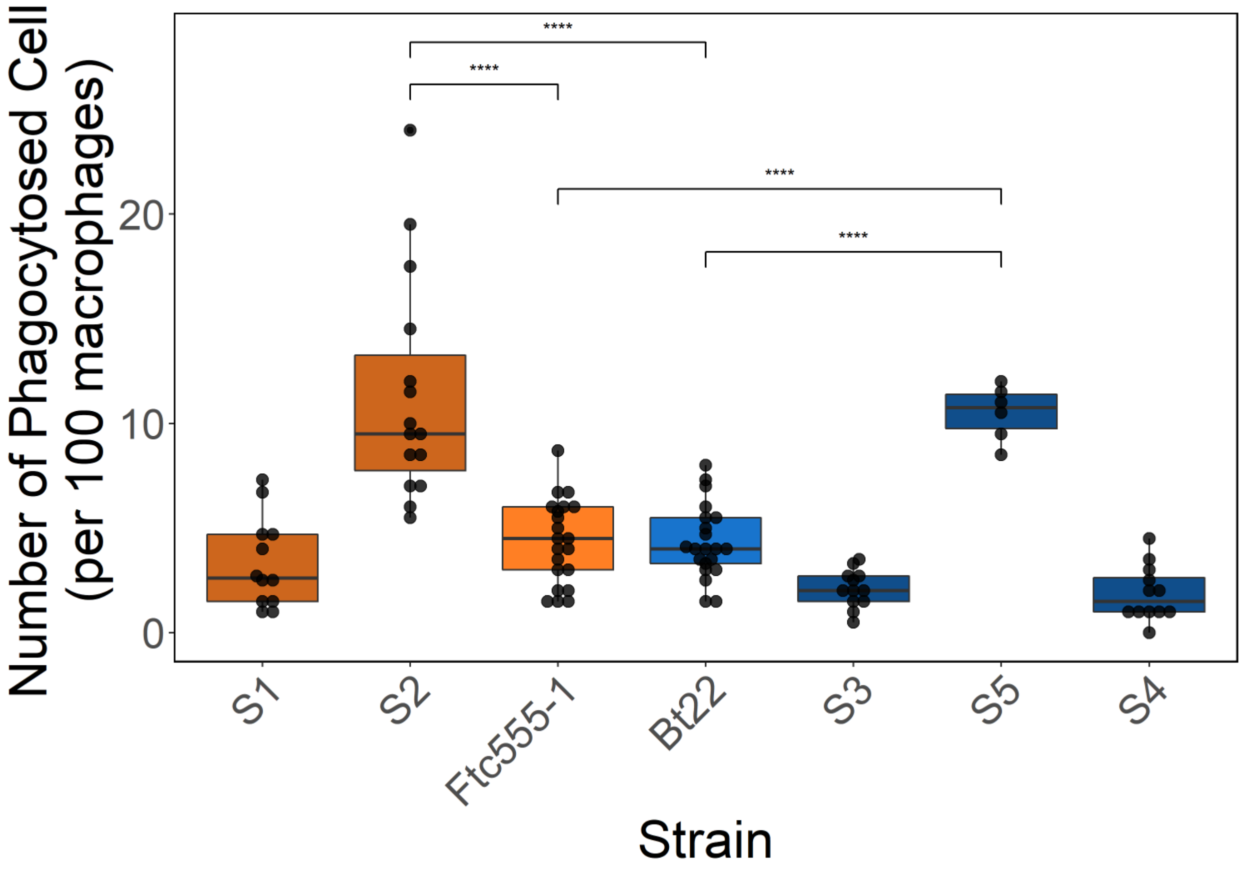

Figure S5: **Phagocytic index is uncorrelated with parental genotype at the chromosome 8 QTL.** Boxplots representing the phagocytic index of parental strains and segregants. Strains are oriented in rank order of amoeba sensitivity. Boxplots are colored by chromosome 8 QTL genotype. Orange indicates strains with the Ftc555-1 allele and blue those with the Bt22 allele. Significance determined by ANOVA ($F = 22.18$; $p<0.0001$).
